# Design of a Multi-Epitope Vaccine using β-barrel Outer Membrane Proteins Identified in *Chlamydia trachomatis*

**DOI:** 10.1101/2025.07.10.664186

**Authors:** Amisha Panda, Jahnvi Kapoor, B. Hareramadas, Ilmas Naqvi, Satish Ganta, Ravindresh Chhabra, Sanjiv Kumar, Anannya Bandyopadhyay

## Abstract

*Chlamydia trachomatis* is an obligate intracellular Gram-negative pathogen responsible for causing sexually transmitted infections (STIs) and trachoma. Current interventions, including screening and antibiotics, are limited due to the widespread nature of asymptomatic infections, and the absence of licensed vaccine exacerbates the challenge. In this study, we predicted outer membrane β-barrel (OMBB) proteins and designed a multi-epitope vaccine (MEV) construct using identified proteins. We employed a consensus-based computational framework on the *C. trachomatis* D/UW-3/CX proteome and identified 17 OMBB proteins, including well-known Pmp family members and MOMP. Eight OMBB proteins were computationally characterized, which showed significant structural homology with known outer membrane proteins from other bacteria. Sequence-based annotation tools were used to determine their putative functions. B-cell and T-cell epitopes were predicted from the selected proteins. The MEV construct was designed using four cytotoxic T lymphocyte (CTL) epitopes and 29 helper T lymphocyte (HTL) epitopes predicted from six OMBB proteins, which were conserved across 106 *C. trachomatis* serovars. The vaccine was supplemented at the N-terminus with Cholera enterotoxin subunit B and PADRE sequence to enhance its immunogenicity. The MEV construct of 780 amino acids was antigenic, non-allergenic, non-toxic, and soluble. Secondary structure analysis revealed 95% random coils. The 3D structural model of MEV was generated and validated, confirming its structural reliability. Molecular docking between MEV and Toll-like receptor 4 (TLR4) revealed strong and stable binding interactions, supporting its potential to elicit a strong immune response. This study highlights OMBB proteins as promising immunogenic targets and presents a computationally designed MEV candidate for *C. trachomatis* infection.

## 1. Introduction

Globally, *Chlamydia trachomatis* is the leading cause of bacterial sexually transmitted infections (STIs) in humans, with approximately 131 million new cases reported annually (Newman et al. 2015; Centers for Disease Control and Prevention 2018). It is a major cause of morbidity with an estimated prevalence of 1–2% in the United States (De Schryver and Meheus 1990; K. Gupta et al. 2021; Rodrigues, Sousa, and Vale 2022). There are 19 major serovars of *C. trachomatis*, classified into three distinct groups, based on tissue tropism and the diseases they cause: ocular trachoma (A, B, Ba, C); urogenital infections (D, Da, E, F, G, Ga, H, I, Ia, J, K), and lymphogranuloma venereum (LVG) disease (L1, L2, L2a, L3) (Lesiak-Markowicz et al. 2019; Morré et al. 2000; Lanfermann et al. 2021; Pilo et al. 2021). *C. trachomatis* is an obligate intracellular, Gram-negative, ovoid-shaped, immobile bacterium that belongs to the family Chlamydiaceae within the order Chlamydiales (Elwell, Mirrashidi, and Engel 2016; Nunes and Gomes 2014). Its unique life cycle alternates between two morphological forms: the infectious, extracellular elementary bodies (EBs) and the intracellular, replicative reticulate bodies (RBs). EBs infect host cells and differentiate into RBs, which replicate within the host cell. After 1–3 days, RBs re-differentiate to EBs, which are released upon cell lysis, initiating the next infection cycle. The bacteria is primarily transmitted through sexual contact but can also be passed from an infected mother to her newborn during vaginal canal delivery (Malhotra et al. 2013). *Chlamydia* primarily infects columnar epithelial cells in the cervix, urethra, rectum, and other non-genital areas (Malik et al. 2006; Fan and Zhong 2015; Robert J. Suchland et al. 2008; R. J. Suchland et al. 2000). Infections caused by this bacterium are often asymptomatic and undiagnosed but persistent or repeated. They can result in severe complications, such as cervicitis, urethritis, pelvic inflammatory disease (PID), cervical cancer, and infertility. *C. trachomatis* can also infect the eye, causing the disease trachoma, which is the leading cause of infectious blindness worldwide (de la Maza, Zhong, and Brunham 2017). The direct lifetime medical costs of *C. trachomatis* infections were estimated to be approximately $691 million in the US alone in 2018 (Chesson et al. 2021). Treatment for known exposures or symptomatic infections involves antibiotics; however, the widespread nature of asymptomatic infections underscores the need for the development of protective vaccines (de la Maza, Darville, and Pal 2021).

*C. trachomatis* D/UW-3/CX is a serovar D strain, originally isolated from the cervix of an asymptomatic patient. It is the reference strain, and has a genome of approximately 1.04 Mb and contains 887 protein-coding genes (Sayers et al. 2023). Like other Gram-negative bacteria, *Chlamydia* species have a cell envelope comprising an outer membrane (OM), inner membrane (IM), and periplasmic space (Tamura et al. 1971). The outer leaflet of the OM contains lipopolysaccharides (Filip et al. 1973). The chlamydial cell envelope is characterized by several unique features: its distinct peptidoglycan structure (Chopra et al. 1998; Ghuysen and Goffin 1999; Moulder 1993); a disulfide-bond cross-linked major outer membrane protein (MOMP) in the OM; and cross linked cysteine-rich proteins (CRPs) within the periplasm (Hatch 1996). Bacteria with diderm envelopes possess a diverse family of outer membrane proteins (OMPs), characterized by β-barrel structures and lipopolysaccharides (Koebnik, Locher, and Van Gelder 2000; Fairman, Noinaj, and Buchanan 2011). β-barrels are cylindrical proteins made up of amphipathic anti-parallel β-strands, typically comprising 8 to 36 β-strands (Horne, Brockwell, and Radford 2020). They are involved in a variety of essential cellular functions, such as nutrient acquisition, membrane biogenesis, OMP assembly, adhesion, biofilm formation, efflux, proteolysis, and pilus formation (Solan et al. 2021). In 1981, Caldwell et al. observed that the hydrophobic OMPs of *Chlamydia* could be isolated from soluble EB proteins using mild detergents like Triton X-114, sarkosyl, and octylglucoside. This sarkosyl-insoluble fraction was referred to as the chlamydial outer membrane complex (COMC) (Caldwell, Kromhout, and Schachter 1981). The COMC, a stabilizing network of disulfide-linked CRPs, provides rigidity to the cell envelope, thus ensuring the survival of the bacterium in harsh extracellular conditions. The COMC is primarily composed of MOMP, several polymorphic membrane proteins (Pmps), and two CRPs: OmcA and OmcB (Hatch, Allan, and Pearce 1984; Stephens et al. 1998; Sardinia, Segal, and Ganem 1988; Liu et al. 2010). Other proteins associated with the COMC include PorB, Omp85, SctC, CTL0887, CTL0541, CTL0645, OprB, and Pal (Liu et al. 2010). Although *C. trachomatis* OMPs (MOMP and Pmps) have been identified and structurally characterized, many outer membrane β-barrel (OMBB) proteins are yet to be identified. Identification and characterization of these novel OMBB proteins would potentially offer valuable insights into the physiology, and virulence of the bacterium, that would be imperative for the development of diagnostics, vaccines, or therapeutic interventions.

Early detection and treatment of chlamydial infections are crucial for reducing the spread of the disease. Although effective antibiotics are available to treat *C. trachomatis* infections, vaccination remains the most affordable and promising strategy (Zhong et al. 2019; Rodrigues et al. 2022). Currently, no licensed human vaccine is available; however, several approaches have been explored for designing vaccines against *C. trachomatis* infection. Initial whole organism-based vaccines (live or inactivated) offered only limited protection against trachoma, leading to short-lived, serovar-specific immune responses and occasional hypersensitivity upon re-exposure (S. P. Wang and Grayston 1967; Taylor 2008). Subsequent second-generation subunit vaccines targeting chlamydial proteins like Pmps (Vasilevsky et al. 2016), Chlamydial Protease-Like Activity Factor (CPAF) (Murthy et al. 2011), and MOMP (Sukumar Pal, Peterson, and de la Maza 2005) were explored preclinically, with MOMP showing strong protective immunity in animal models (Farris and Morrison 2011; Kari et al. 2009; Simpson et al. 2023; Olsen et al. 2022). However, third-generation DNA-based vaccines were found to be ineffective (S. Pal et al. 1999; L. Wang et al. 2017). Currently, the CTH522 vaccine, formulated with CAF®01 adjuvant, has successfully completed a Phase I clinical trial. This vaccine specifically targets the variable domain 4 (VD4) region of MOMP-a well-characterized β-barrel protein of *C. trachomatis* (Olsen et al. 2021). This highlights the importance of OMBB proteins for designing peptide-based multi-epitope vaccines, which could offer a more effective solution to control the disease.

In the present study, we predicted OMBB proteins and designed a multi-epitope vaccine (MEV) construct using identified proteins. Using a combination of tools that predict OM and β-barrel proteins, we predicted eight putative OMBB proteins. Structural homologs and potential functions were predicted using AlphaFold 3 generated structural models. Sequence-based functional annotation was performed to predict the potential functions of OMBB proteins. Moreover, sequence variations from 106 *C. trachomatis* strains were identified and mapped onto the structural models, revealing that many variations occurred in surface-exposed loops, indicative of selective pressure. To harness the immunogenic potential of these proteins, we predicted B-cell, cytotoxic T-lymphocyte (CTL), and helper T-lymphocyte (HTL) epitopes. Surface-exposed MHC-class I and MHC-class II epitopes from six OMBB proteins were linked together using AAY and GPGPG linkers, respectively to design the MEV construct. The physicochemical properties of the MEV were analyzed, and its secondary and tertiary structures were predicted. Molecular docking between the MEV and TLR4 was performed to assess the binding interaction. This study underscores the potential of OMBB proteins as targets for developing an MEV against *C. trachomatis* infection, offering a promising framework to tackle this global pathogen.

## 2. Materials and Methods

### 2.1 Prediction of outer membrane β-barrel proteins

We selected *C. trachomatis* D/UW-3/CX, the reference strain for our study. Using genome assembly ASM872v1, amino acid sequences of all 887 proteins of *C. trachomatis* D/UW-3/CX proteome were downloaded from NCBI (Accessed on 12 June, 2024) (Sayers et al. 2023).

Pepstats tool of the EMBOSS package was used on all protein sequences to determine the peptide length, molecular weight, charge, and isoelectric point (pI) (Rice, Longden, and Bleasby 2000). SPAAN (Accessed on 2 July, 2024) was utilized to predict the adhesins or adhesin like proteins (Sachdeva et al. 2005). SignalP 5.0 (Accessed on 13 June, 2024) was used to predict signal peptide and its cleavage position in the protein sequences (Almagro Armenteros et al. 2019). Subcellular localization of the proteins was predicted using CELLO v.2.5 (Accessed on 22 June, 2024) (C.-S. Yu, Lin, and Hwang 2004) and PSORTb v3.0 (Accessed on 15 June, 2024) (N. Y. Yu et al. 2010). CDD (Conserved Domain Database) (Accessed on 25 September, 2024) search was done to identify the functional domains present within protein sequences (Yang et al. 2020). The computational pipeline designed to select OMPs is schematically represented in Figure 1.

**Figure 1:**
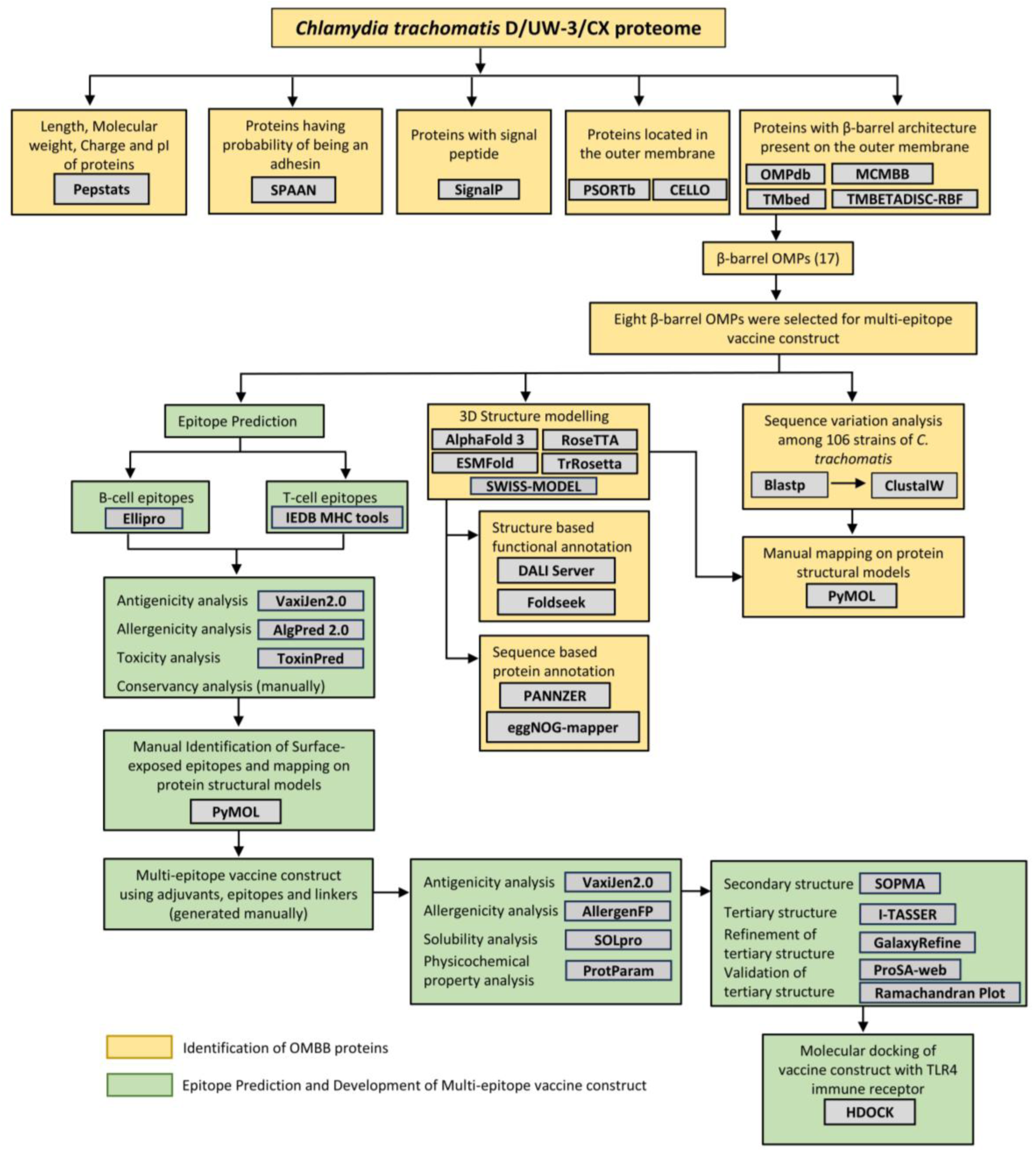
Computational framework for predicting OM-localized, β-barrel proteins and designing of a multi-epitope vaccine against *Chlamydia trachomatis* infection. *C. trachomatis* proteome was mined in silico using various tools such as SignalP, PSORTb, CELLO, OMPdb, MCMBB, TMbed, TMBETADISC-RBF. Size, charge, and isoelectric point of proteins were predicted using Pepstats. Structural models were generated using five modelling tools as shown in the schematic above. Structural models generated using AlphaFold 3 server were used as queries in the DALI and Foldseek servers, and putative functions were annotated. Further, sequence-based protein annotation was done using tools: PANNZER and eggNOG-mapper. Amino acid sequence variation analysis from 106 strains of *C. trachomatis* was performed using ClustalW. B-cell and T-cell epitopes were predicted using Ellipro and IEDB MHC tools, respectively. Antigenicity, allergenicity, and toxicity of predicted epitopes were evaluated using VaxiJen 2.0, AlgPred 2.0, and ToxinPred, respectively. Further, multi-epitope vaccine was constructed using adjuvants, surface-exposed epitopes and linkers. Secondary structure of vaccine construct was predicted using SOPMA tool. Tertiary structure of MEV was generated using I-TASSER tool, followed by refinement of the tertiary structure using GalaxyRefine web server. ProSA-web server and Ramachandran plot were used to evaluate the overall quality of the predicted tertiary structure. Further, HDOCK was used for molecular docking of MEV construct with TLR4 immune receptor (PDB ID: 3FXI).

A consensus-based computational approach was employed to identify OMBB proteins, where the output from four OMP prediction tools (one database-OMPdb (Accessed on 17 June, 2024) (Tsirigos, Bagos, and Hamodrakas 2011), and three tools-MCMBB (Accessed on 23 September, 2024) (Bagos et al. 2004), TMBETADISC-RBF (Accessed on 27 June, 2024) (Ou et al. 2008), and TMbed (Accessed on 27 June, 2024) (Bernhofer and Rost 2022)) was taken into consideration for identification of OMPs. Furthermore, proteins were classified based on the number of tools providing positive results for a given protein sequence. This implied that higher confidence was assigned to those predicted as OMBB protein by more tools.

### 2.2 Structural modelling of predicted outer membrane β-barrel proteins

Structural models of the predicted proteins were obtained from the AlphaFold database (Accessed on 4 October, 2024) (Varadi et al. 2022). To validate the structural models available in the AlphaFold database, we generated 3D structural models using additional four modelling tools: ESMFold (Accessed on 28 December, 2024) (Lin et al. 2023), SWISS-MODEL (Accessed on 20 December, 2024) (A. Waterhouse et al. 2018), RoseTTAFold (Accessed on 24 December, 2024) (Baek et al. 2021) and TrRosetta (Accessed on 24 December, 2024) (Du et al. 2021). For each protein, the top ranked predictions, based on highest confidence scores, were selected for visualization and further analysis. The resulting atomic coordinate files were visualized using PyMOL (“Schrödinger, L., & DeLano, W. (2020). PyMOL” n.d.). To assess structural similarities and differences across models, structural models were aligned using US-align (Accessed on 11 January, 2025) (C. Zhang et al. 2022) online web server. Alignments were visualized using PyMOL, and corresponding figures were generated.

### 2.3 Functional annotation of predicted outer membrane β-barrel proteins

Given that two out of eight OMBB proteins were classified as hypothetical proteins, a structure-based approach was employed to unravel their putative functions. The atomic coordinates of the structural models were submitted as queries to the DALI server (Accessed on 5 October, 2024) (Holm 2020) with the full PDB search option, allowing comparisons against all protein structures available in the Protein Data Bank (PDB). For each query, the protein with the highest Z-score was selected as the top-hit and used as the basis for functional annotation. Functions of the top-hit proteins were searched from the available literature to annotate putative functions to the predicted OMBB proteins. To complement this approach, Foldseek tool (Accessed on 8 April, 2025) (van Kempen et al. 2024) was employed to identify structural homologs across five protein databases: PDB100, CATH50, AFDB50, AFDB-SWISSPROT and AFDB-Proteome. The top hit, based on the highest TM-score across the five protein databases, was selected for functional annotation. In parallel, sequence-based annotation was performed using two tools-PANNZER (Accessed on 6 April, 2025) (Protein ANNotation with Z-scoRE) (Törönen and Holm 2022) and eggNOG-mapper (Accessed on 8 April, 2025) (Cantalapiedra et al. 2021).

### 2.4 Amino acid sequence variation analysis across *C. trachomatis* strains

Predicted OMBB proteins from *C. trachomatis* reference genome D/UW-3/CX were compared across 106 complete *C. trachomatis* genomes, representing various serovars, to study amino acid sequence variation (Table S3). Orthologous sequences for each protein were identified using BLASTP (E-value < 1.0E-03, bitscore > 100) and aligned using ClustalW for Multiple Sequence Alignment (MSA). MSA analysis identified amino acid variations among the orthologs, which were subsequently mapped onto structural models using PyMOL.

### 2.5 B-cell epitope prediction

ElliPro, a web server of the Immune Epitope Database (IEDB) (Accessed on 14 October, 2024), was utilized to predict both linear and conformational B-cell epitopes using the 3D structural model of the proteins. Predictions were performed using default parameters (Ponomarenko et al. 2008).

### 2.6 T-cell epitope prediction

Cytotoxic T lymphocyte (CTL) and helper T lymphocyte (HTL) epitopes were predicted using MHC I and MHC II prediction tools, respectively, of the IEDB (Nielsen, Lundegaard, and Lund 2007). CTL epitope prediction targeted 9-mer peptides with a percentile rank threshold of <0.5. The analysis included the following Human Leukocyte Antigen (HLA) alleles: HLA-A*01:01, HLA-A*02:01, HLA-A*11:01, HLA-A*24:02, HLA-B*07:02, HLA-B*08:01, HLA-B*15:01, HLA-B*44:02, HLA-C*04:01, HLA-C*07:02.

HTL epitopes (15-mer peptides) were predicted with a percentile rank threshold <2 for the following HLA alleles: HLA-DRB1*01:01, HLA-DRB1*03:01, HLA-DRB1*04:01, HLA-DRB1*07:01, HLA-DRB1*08:02, HLA-DRB1*13:01, HLA-DRB1*13:02, HLA-DRB1*15:01.

### 2.7 Homology of predicted epitopes with human proteome

Proteins that are non-homologous to human sequences are less likely to induce an autoimmune response. To address this issue, a comparison was made between Homo sapiens proteome (TaxID: 9606) and predicted peptides using the NCBI BLASTP database (Accessed on 14 October, 2024) (Altschul et al. 1990). Peptides with an E-value exceeding 0.05 were considered non-homologous and suitable candidates for vaccine development.

### 2.8 Antigenicity, allergenicity and toxicity evaluation of predicted epitopes

After screening the predicted epitopes for homology with human proteome, the selected epitopes were evaluated for antigenicity, allergenicity and toxicity. Antigenicity was evaluated using the VaxiJen2.0 online web server (Accessed on 25 November, 2024), employing the bacterial prediction model and a threshold score of 0.4 (Doytchinova and Flower 2007). Allergenicity of the predicted epitopes was assessed using AlgPred 2.0 (Accessed on 25 November, 2024), with default parameters (Sharma et al. 2021). Toxicity of the selected epitopes was evaluated using ToxinPred server (Accessed on 25 November, 2024) (S. Gupta et al. 2013). Epitopes that were identified as antigenic, non-toxic, and non-allergenic were selected for further analysis.

### 2.9 Conservation analysis of predicted epitopes across *C. trachomatis* strains

MSA was performed to assess the conservation of selected epitopes across 106 strains of *C. trachomatis*. Conserved epitopes across 106 *C. trachomatis* strains were selected for further analysis to ensure the broad applicability and effectiveness of the predicted epitopes.

### 2.10 Structural mapping of the predicted epitopes

Epitopes were mapped to determine their location on the structural models of the predicted OMBB proteins. Since a prospective vaccine candidate should be a surface-exposed protein that can stimulate the host immune system for clearance of the pathogen, epitopes located on the extracellular loop (ECL) region were considered for further analysis.

### 2.11 Development of multi-epitope vaccine construct

Antigenic, non-allergenic, non-toxic, conserved, and surface-exposed epitopes were selected for the construction of MEV. To construct the vaccine sequence, epitopes were joined together with appropriate peptide linkers (Dong et al. 2020). HTL epitopes were connected using GPGPG linkers, while CTL epitopes were connected with AAY linkers. To enhance immunogenicity, two adjuvants-Cholera toxin B subunit (CTB) (accession no: P01556) and PADRE sequence were added at the N-terminus with the help of EAAAK linkers. A polyhistidine tag for purification was added at the C-terminus through a KK linker to obtain the complete vaccine protein sequence (Z. Chen et al. 2021).

### 2.12 Evaluation of multi-epitope vaccine construct

The MEV was evaluated using bioinformatics tools. ProtParam (Accessed on 22 January, 2025) was used to analyze physicochemical properties (Wilkins et al. 1999), VaxiJen 2.0 for antigenicity assessment (Doytchinova and Flower 2007), SOLpro (Accessed on 18 February, 2025) for solubility prediction (Magnan, Randall, and Baldi 2009) and AllergenFP v1.1 (Accessed on 22 January, 2025) for allergenicity evaluation (Dimitrov et al. 2014).

### 2.13 Structure prediction of the vaccine construct and assessment of model quality

SOPMA secondary structure prediction tool (Accessed on 22 March, 2025) was used to predict the secondary structures of MEV (Geourjon and Deléage 1995). SOPMA analyzes and assesses the structure of the target protein by predicting the formation of α-helices, beta-turns, random coils, and extended strands. Tertiary structure of MEV was generated using I-TASSER tool (Accessed on 25 March, 2025) (Y. Zhang 2008), followed by refinement of the tertiary structure using GalaxyRefine web server (Accessed on 26 March, 2025) to improve model quality (Heo, Park, and Seok 2013). The quality of the MEV tertiary structure model was assessed using the SWISS-MODEL Structure Assessment tool (Accessed on 5 April, 2025) (A. M. Waterhouse et al. 2024) and the ProSA-web server (Accessed on 6 April, 2025) (Wiederstein and Sippl 2007). Model validation was based on Z-score and the distribution of amino-acid residues in the Ramachandran plot, providing insights into the overall stability and stereochemical quality of the predicted structure, respectively.

### 2.14 Molecular docking of vaccine construct with TLR4 immune receptor

To evaluate the interaction between the MEV construct and host immune receptors, molecular docking was performed with Toll-like receptor 4 (TLR4), which plays a key role in recognizing *Chlamydia* and initiating immune responses (Viana Invenção et al. 2021; Nosratababadi et al. 2017). The atomic coordinate file of TLR4 (PDB ID: 3FXI) was obtained from RCSB PDB database (Accessed on 10 April, 2025). Since the structure was co-crystallized with *E. coli* lipopolysaccharide (LPS) molecules, we cleaned the atomic coordinate file by removing LPS molecules. Docking was performed using the HDOCK server (Accessed on 10 April, 2025). The resulting MEV-TLR4 complex was analyzed for interacting amino acid residues using PDBsum server and the docked complex was visualized using PyMOL.

## 3. Results

### 3.1 Outer membrane β-barrel protein prediction

A consensus-based computational framework was employed to identify OMBB proteins in the *C. trachomatis* D/UW-3/CX genome, which encodes 887 proteins (Figure 1). The framework included nine computational tools: Pepstats, SPAAN, SignalP, CELLO, PSORTb, OMPdb, MCMBB, TMBETADISC-RBF, and TMbed. Prediction outputs from all the tools were integrated for each protein (Table S1). OMBB-specific predictions were derived from OMPdb, MCMBB, TMBETADISC-RBF, and TMbed (Table 1). Through consensus-based screening criteria and manual curation based on structural models, seventeen OMBB proteins were identified. Since nine Pmps (PmpA, PmpB, PmpC, PmpD, PmpE, PmpF, PmpG, PmpH, and PmpI) have been identified (Table S1) and structurally well-characterized (Cervantes et al. 2024), eight OMBB proteins were selected for further analysis. These eight proteins were subsequently classified into three groups based on the level of consensus among the OMBB prediction tools: Group A, consisted of one protein (NP_219746.1) predicted as OMBB by all four tools, Group B included three proteins (NP_220140.1, NP_220200.1 and NP_220232.1) predicted by any three tools; and Group C comprised four proteins (NP_219509.1, NP_219858.1, NP_219881.1, and NP_219899.1) predicted as OMBB by any two tools (Table 1). Structural models were generated using five different modelling tools: AlphaFold 3 (Abramson et al. 2024), ESMFold, SWISS-MODEL, RoseTTAFold, and TrRosetta. To quantitatively assess the structural similarity, structural models were aligned using the US-Align server, and the resulting RMSD values were found to be low (Table S2).

**Table 1.**
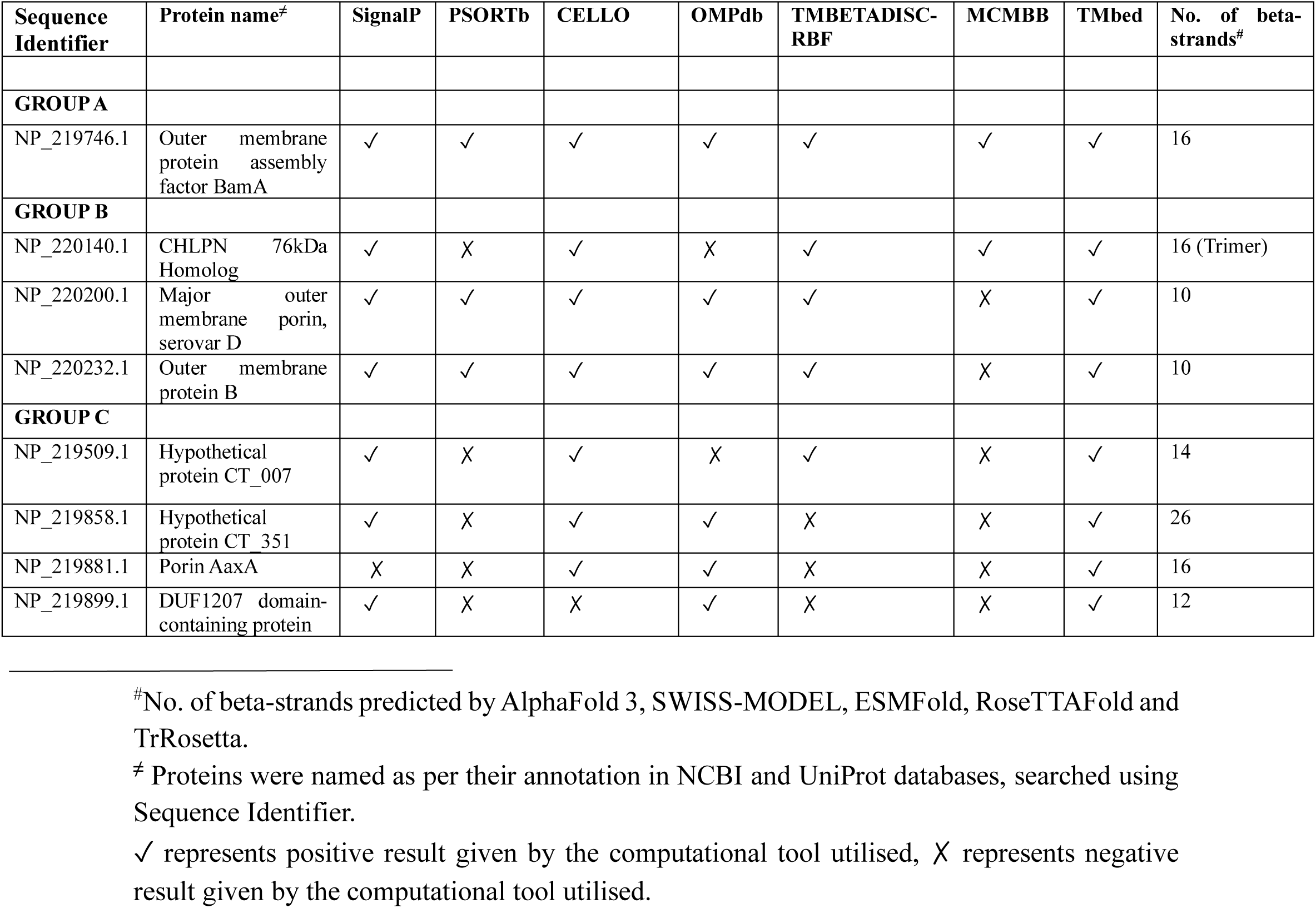
List of eight OMBB proteins selected for multi-epitope vaccine construct using consensus-based computational approach.

| Sequence Identifier | Protein name<sup>‡</sup> | SignalP | PSORTb | CELLO | OMPdb | TMBETADISC-RBF | MCMBB | TMbed | No. of beta-strands<sup>#</sup> |
| --- | --- | --- | --- | --- | --- | --- | --- | --- | --- |
| **GROUP A** |  |  |  |  |  |  |  |  |  |
| NP_219746.1 | Outer membrane protein assembly factor BamA | ✓ | ✓ | ✓ | ✓ | ✓ | ✓ | ✓ | 16 |
| **GROUP B** |  |  |  |  |  |  |  |  |  |
| NP_220140.1 | CHLPN 76kDa Homolog | ✓ | X | ✓ | X | ✓ | ✓ | ✓ | 16 (Trimer) |
| NP_220200.1 | Major outer membrane porin, serovar D | ✓ | ✓ | ✓ | ✓ | ✓ | X | ✓ | 10 |
| NP_220232.1 | Outer membrane protein B | ✓ | ✓ | ✓ | ✓ | ✓ | X | ✓ | 10 |
| **GROUP C** |  |  |  |  |  |  |  |  |  |
| NP_219509.1 | Hypothetical protein CT_007 | ✓ | X | ✓ | X | ✓ | X | ✓ | 14 |
| NP_219858.1 | Hypothetical protein CT_351 | ✓ | X | ✓ | ✓ | X | X | ✓ | 26 |
| NP_219881.1 | Porin AaxA | X | X | ✓ | ✓ | X | X | ✓ | 16 |
| NP_219899.1 | DUF1207 domain-containing protein | ✓ | X | X | ✓ | X | X | ✓ | 12 |

<sup>#</sup>No. of beta-strands predicted by AlphaFold 3, SWISS-MODEL, ESMFold, RoseTTAFold and TrRosetta.
<sup>‡</sup> Proteins were named as per their annotation in NCBI and UniProt databases, searched using Sequence Identifier.
✓ represents positive result given by the computational tool utilised, X represents negative result given by the computational tool utilised.

#### 3.1.1 Group A

**Group A** comprised the protein, NP_219746.1, which is annotated as the outer membrane assembly factor BamA in the UniProt database (Table 1). BamA functions as the core component of the β-barrel assembly machine (BAM) complex, which, along with BamBCDE, is responsible for the insertion and assembly of β-barrel proteins into the OM (Bakelar, Buchanan, and Noinaj 2016). The structural model of NP_219746.1 revealed a characteristic bipartite structure of BamA, consisting of a periplasmic N-terminal region and a C-terminal 16-stranded β-barrel domain (Figure 2). The N-terminal segment contains five characteristic polypeptide transport-associated (POTRA) domains (P1–P5), which serve as a scaffold for the binding of BamBCDE proteins and facilitate OMBB protein folding (Konovalova, Kahne, and Silhavy 2017). CDD search confirmed its homology to BamA (Table S1). Additionally, structural alignment tool (DALI) and sequence-based annotation tools (eggNOG-mapper and PANNZER) consistently identified NP_219746.1 as a BamA homolog; however, Foldseek tool showed the best structural match with an uncharacterized protein of *Parachlamydia acanthamoebae* UV-7 (Table 2).

**Figure 2:**
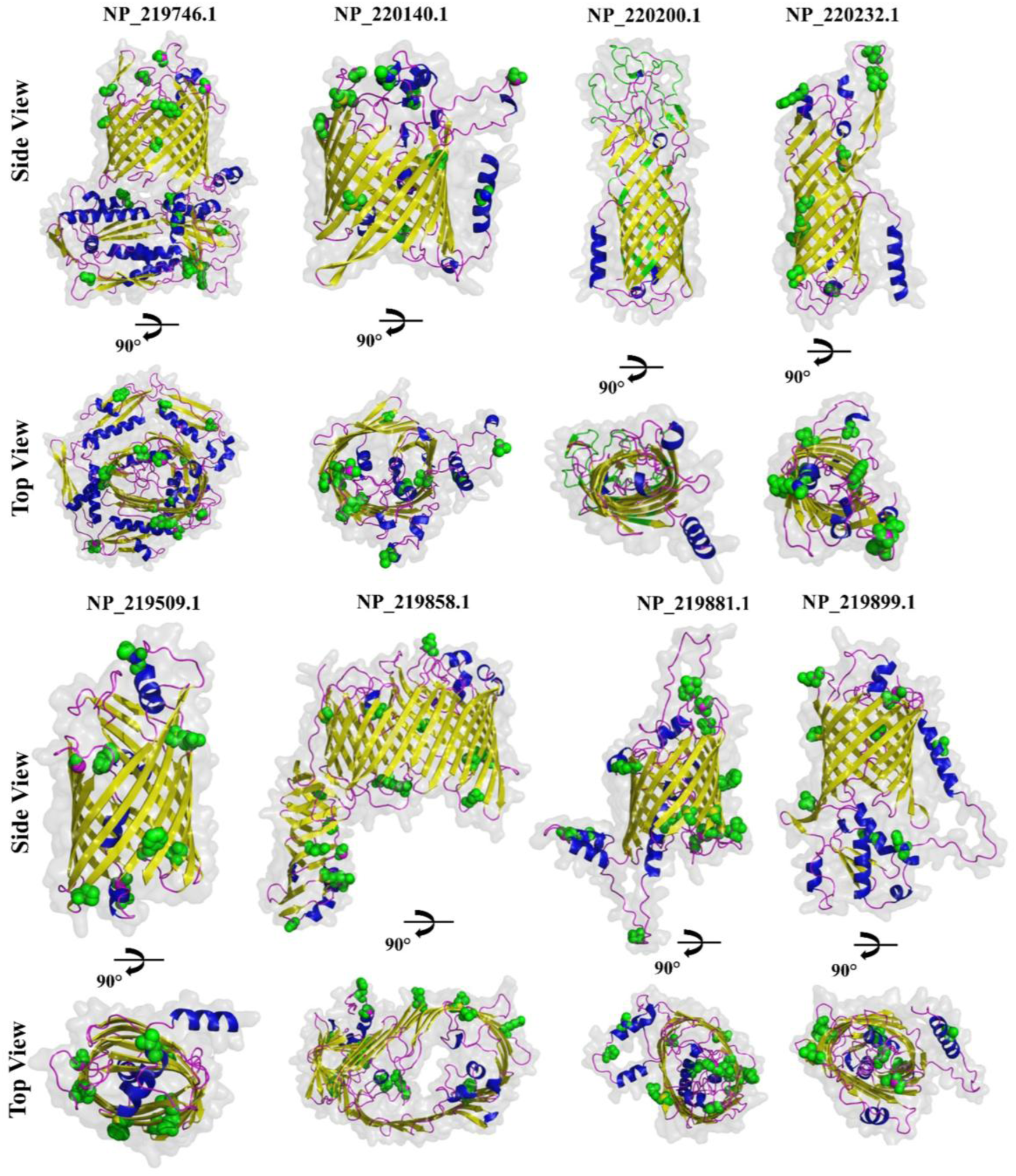
Structural models of β-barrel outer membrane proteins. Structural models of 26-stranded β-barrel protein (NP_219858.1); 16-stranded β-barrel proteins (NP_219746.1, NP_220140.1, and NP_219881.1); 14-stranded β-barrel protein (NP_219509.1); 12-stranded β-barrel protein (NP_219899.1), and 10-stranded β-barrel proteins (NP_220200.1 and NP_220232.1) were generated using AlphaFold 3 server. β-strands, α-helices, and loops are shown in yellow, blue, and magenta colors, respectively. Green spheres indicate amino acid variations across 106 *C. trachomatis* strains. Since NP_220200.1 exhibits amino acid sequence variations at 93 positions, therefore the variations are shown using ribbon representation in green colour.

**Table 2.**
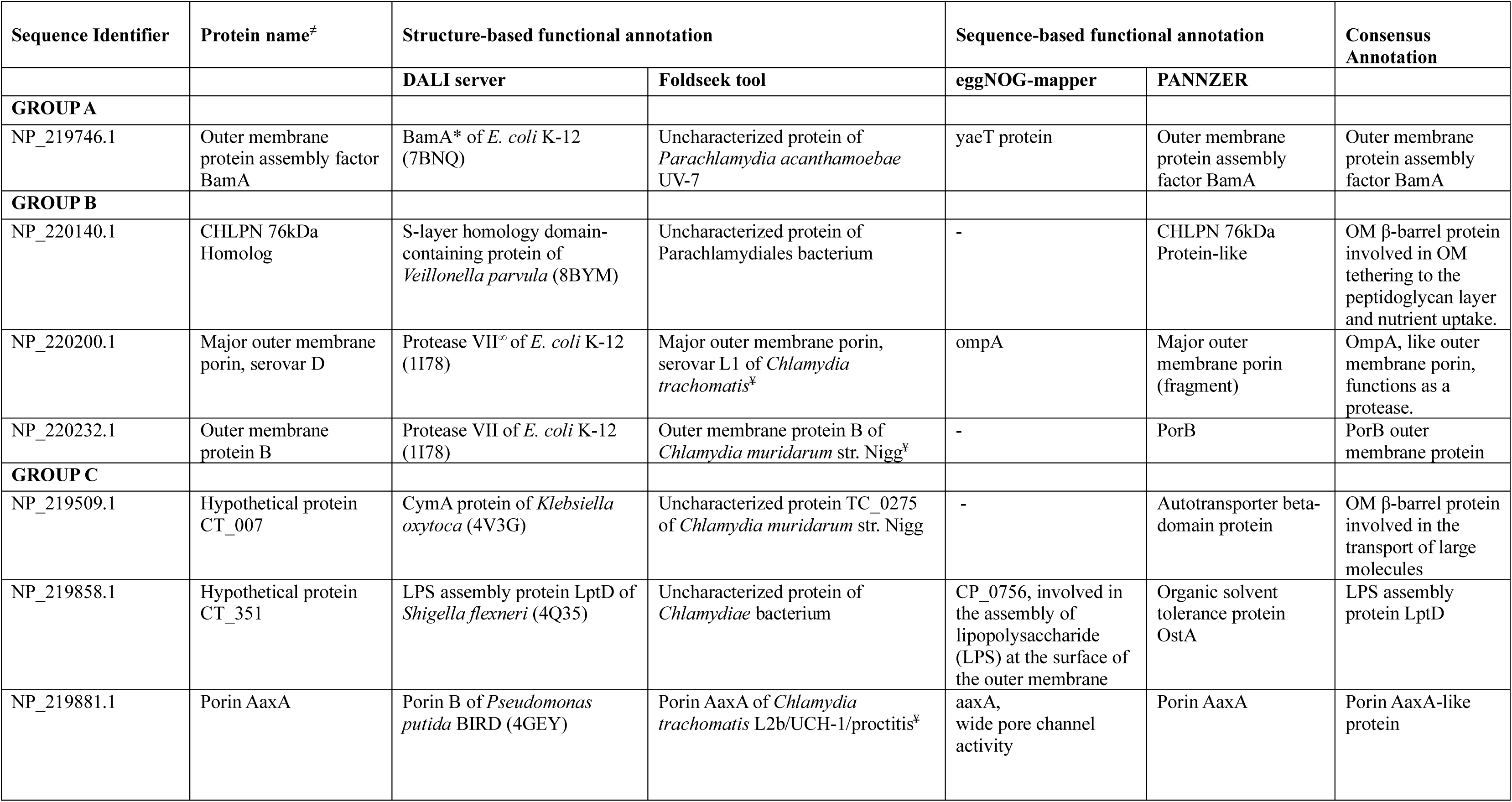
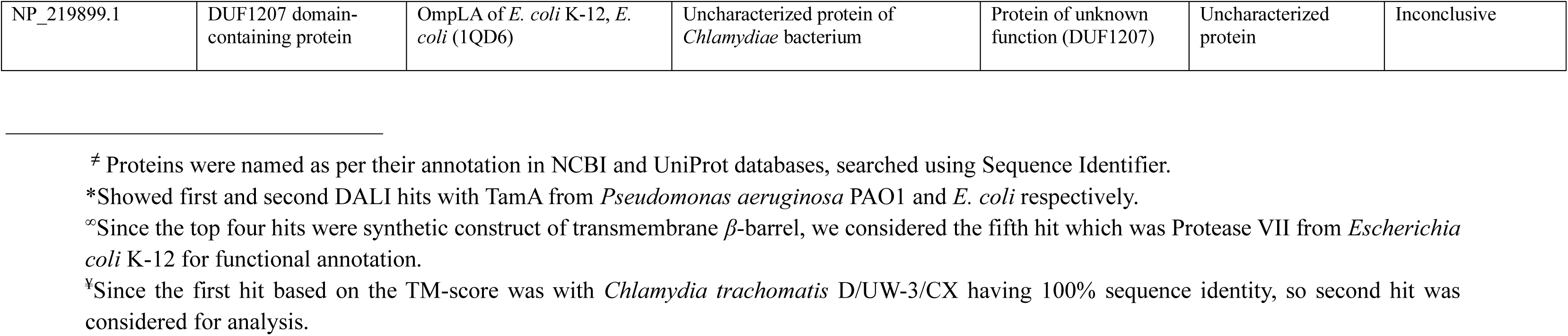
Structure-based and sequence-based functional annotation.

| Sequence Identifier | Protein name<sup>‡</sup> | Structure-based functional annotation |  | Sequence-based functional annotation |  | Consensus Annotation |
| --- | --- | --- | --- | --- | --- | --- |
|  |  | DALI server | Foldseek tool | eggNOG-mapper | PANNZER |  |
| **GROUP A** |  |  |  |  |  |  |
| NP_219746.1 | Outer membrane protein assembly factor BamA | BamA* of *E. coli* K-12 (7BNQ) | Uncharacterized protein of *Parachlamydia acanthamoebae* UV-7 | yaeT protein | Outer membrane protein assembly factor BamA | Outer membrane protein assembly factor BamA |
| **GROUP B** |  |  |  |  |  |  |
| NP_220140.1 | CHLPN 76kDa Homolog | S-layer homology domain-containing protein of *Veillonella parvula* (8BYM) | Uncharacterized protein of Parachlamydiales bacterium | - | CHLPN 76kDa Protein-like | OM β-barrel protein involved in OM tethering to the peptidoglycan layer and nutrient uptake. |
| NP_220200.1 | Major outer membrane porin, serovar D | Protease VII<sup>∞</sup> of *E. coli* K-12 (1I78) | Major outer membrane porin, serovar L1 of *Chlamydia trachomatis*<sup>‡</sup> | ompA | Major outer membrane porin (fragment) | OmpA, like outer membrane porin, functions as a protease. |
| NP_220232.1 | Outer membrane protein B | Protease VII of *E. coli* K-12 (1I78) | Outer membrane protein B of *Chlamydia muridarum* str. Nigg<sup>‡</sup> | - | PorB | PorB outer membrane protein |
| **GROUP C** |  |  |  |  |  |  |
| NP_219509.1 | Hypothetical protein CT_007 | CymA protein of *Klebsiella oxytoca* (4V3G) | Uncharacterized protein TC_0275 of *Chlamydia muridarum* str. Nigg | - | Autotransporter beta-domain protein | OM β-barrel protein involved in the transport of large molecules |
| NP_219858.1 | Hypothetical protein CT_351 | LPS assembly protein LptD of *Shigella flexneri* (4Q35) | Uncharacterized protein of *Chlamydiae* bacterium | CP_0756, involved in the assembly of lipopolysaccharide (LPS) at the surface of the outer membrane | Organic solvent tolerance protein OstA | LPS assembly protein LptD |
| NP_219881.1 | Porin AaxA | Porin B of *Pseudomonas putida* BIRD (4GEY) | Porin AaxA of *Chlamydia trachomatis* L2b/UCH-1/proctitis<sup>‡</sup> | aaxA, wide pore channel activity | Porin AaxA | Porin AaxA-like protein |
| NP_219899.1 | DUF1207 domain-containing protein | OmpLA of *E. coli* K-12, *E. coli* (1QD6) | Uncharacterized protein of *Chlamydiae* bacterium | Protein of unknown function (DUF1207) | Uncharacterized protein | Inconclusive |

<sup>‡</sup> Proteins were named as per their annotation in NCBI and UniProt databases, searched using Sequence Identifier.
\*Showed first and second DALI hits with TamA from *Pseudomonas aeruginosa* PAO1 and *E. coli* respectively.
<sup>∞</sup>Since the top four hits were synthetic construct of transmembrane β-barrel, we considered the fifth hit which was Protease VII from *Escherichia coli* K-12 for functional annotation.
<sup>‡</sup>Since the first hit based on the TM-score was with *Chlamydia trachomatis* D/UW-3/CX having 100% sequence identity, so second hit was considered for analysis.

**Table 3.** Predicted surface-exposed MHC Class-I epitopes for the identified OMBBs.

| Sequence Identifier | Protein name<sup>‡</sup> | HLA allele | Start Position | End Position | Peptide | Score |
| --- | --- | --- | --- | --- | --- | --- |
| NP_219509.1 | Hypothetical protein CT_007 | HLA-C*04:01 | 253 | 261 | IFEYSGREI | 0.248849 |
| NP_219858.1 | Hypothetical protein CT_351 | HLA-A*11:01 | 339 | 347 | KVNSFQNVK | 0.896197 |
|  |  | HLA-B*08:01 | 305 | 313 | DIFPNNFSL | 0.264336 |
| NP_219881.1 | Porin AaxA | HLA-A*11:01 | 273 | 281 | SIKWSNLTK | 0.773741 |

<sup>‡</sup> Proteins were named as per their annotation in NCBI and UniProt databases, searched using Sequence Identifier.

**Table 4.**
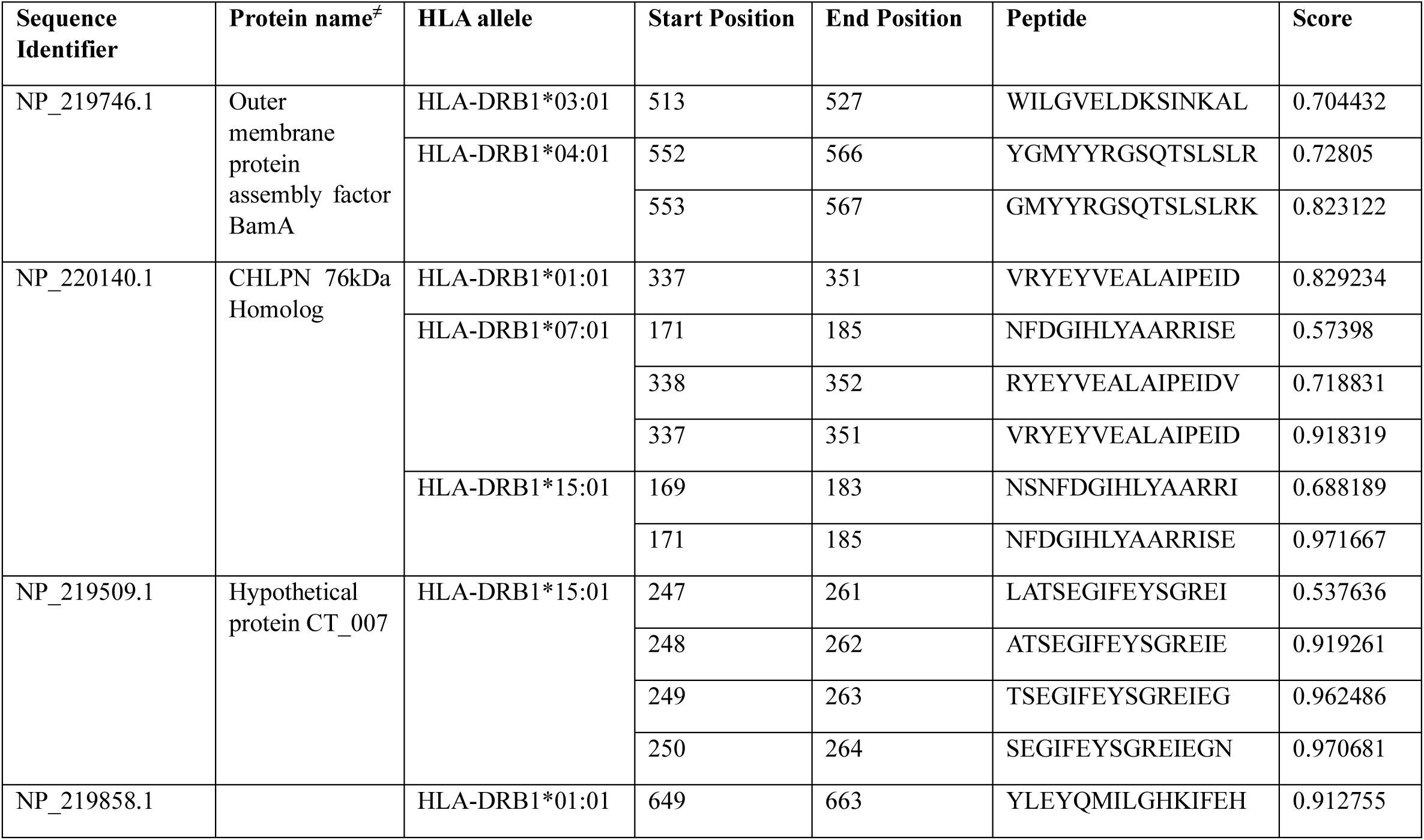
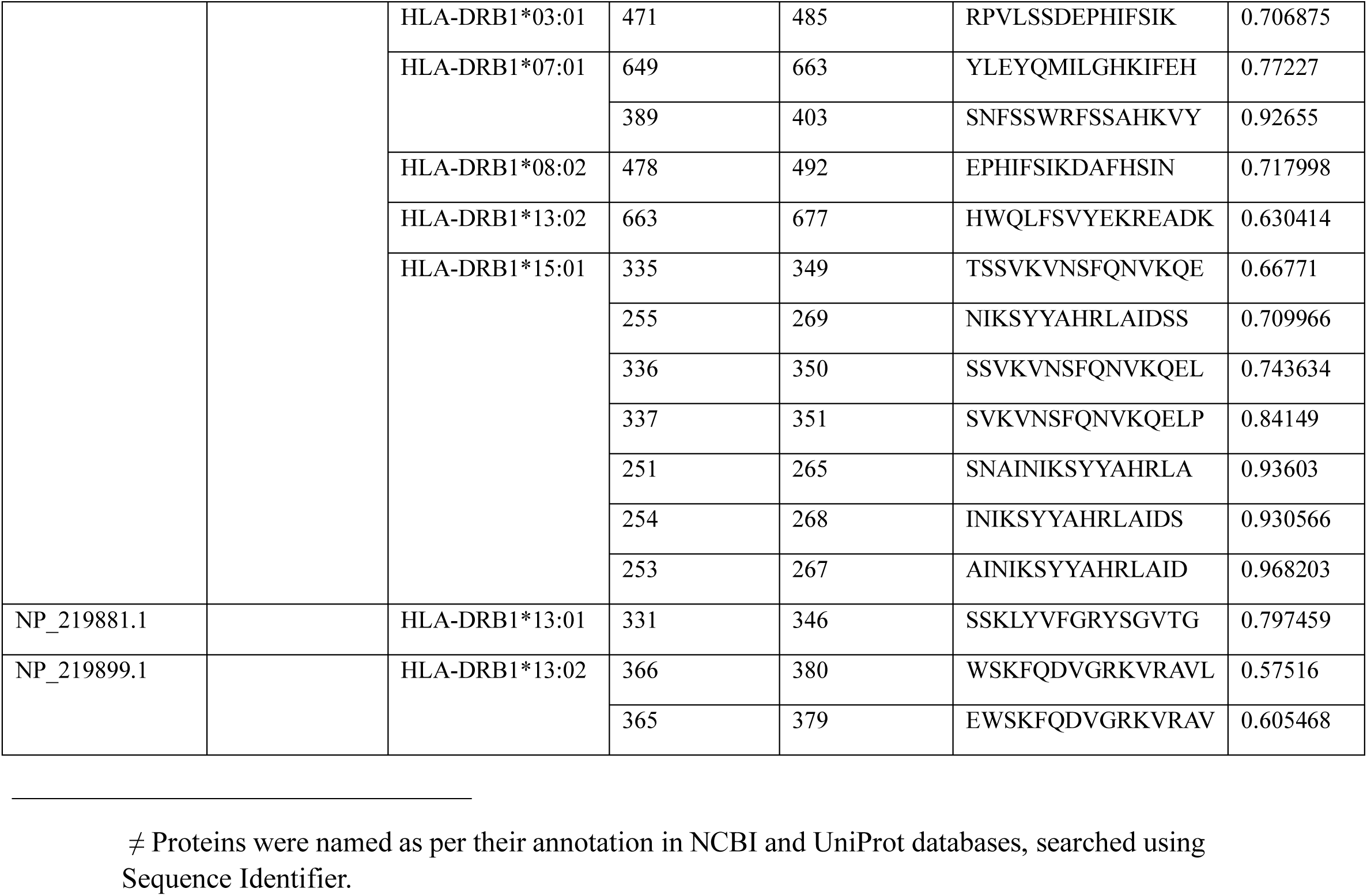
Predicted surface-exposed MHC Class-II epitopes for the identified OMBBs.

#### 3.1.2 Group B

**Group B** comprised three proteins: NP_220140.1, NP_220200.1 (MOMP), and NP_220232.1. Structural models revealed that NP_220140.1 contains a 16-stranded β-barrel domain, while NP_220200.1 and NP_220232.1 each comprise a ten-stranded β-barrel domain (Figure 2).

NP_220140.1 is annotated as ‘CHLPN 76kDa Homolog’ in UniProt; however, it is annotated as a ‘hypothetical protein CT_623’ in NCBI. Structure-based functional annotation using DALI server identified structural homology with outer membrane attachment porin OmpM1 from *Veillonella parvula* (PDB ID: 8BYM) (Table 2), a trimeric β-barrel involved in OM tethering to the peptidoglycan layer and nutrient uptake (Silale et al. 2023). The trimeric model, generated using AlphaFold 3, exhibited strong structural confidence (pTM = 0.78, ipTM = 0.76), suggesting a potential trimeric structure (Figure S1). Foldseek aligned the structure with an uncharacterized protein from a *Parachlamydiales* bacterium, and PANNZER annotated it as ‘CHLPN 76kDa Protein-like’ (Table 2). Combining these annotations, it can be speculated that it is a β-barrel protein involved in OM tethering to the peptidoglycan layer and nutrient uptake.

NP_220200.1, annotated as major outer membrane porin (MOMP) in both NCBI and UniProt, is encoded by the ompA gene. It is highly abundant, contributing approximately 60% of the total COMC (Caldwell, Kromhout, and Schachter 1981). MOMP is 40 kDa in size (G. E. Schulz 2000; Georg E. Schulz 2002; Nikaido 2003) that predominantly forms a trimeric β-barrel architecture, creating ∼2 nm wide pore (Sun et al. 2007). It is a highly antigenic protein and a primary target for vaccine development (de la Maza, Darville, and Pal 2021; Su et al. 1990). *C. trachomatis* MOMP comprises four variable domains (VD) interspersed with five constant domains. Structural modelling revealed a ten-stranded β-barrel architecture and a large unstructured region on the extracellular side (Figure 2). The trimeric model, generated using AlphaFold 3, exhibited strong structural confidence (pTM = 0.8, ipTM = 0.82), suggesting a potential trimeric structure. Although Feher et al. modelled MOMP as a 16-stranded β-barrel based on earlier topology predictions (Feher et al. 2013), structural models generated using AlphaFold3, TrRosetta, RoseTTAFold, ESMFold, and SWISS-MODEL consistently revealed a ten-stranded β-barrel architecture. This suggests that MOMP may adopt alternative β-barrel topologies, highlighting the need for further experimental validation. NP_220200.1 showed the best structural alignment with protease OmpT of *E. coli* K-12 (PDB ID: 1I78) (Table 2) using the DALI server, while Foldseek showed the best structural match with MOMP of *C. trachomatis* serovar L1. Complementing this, eggNOG-mapper and PANNZER associate NP_220200.1 with the major outer membrane porin (OmpA). Together, these findings suggest that NP_220200.1, the major outer membrane porin (OmpA), may function as a protease (Table 2).

NP_220232.1 is annotated as outer membrane protein B (PorB) in NCBI and UniProt databases. Structural modelling revealed a ten-stranded β-barrel architecture with a large unstructured region (∼112 amino acid residues) on the extracellular side (Figure 2). Structural alignment using DALI server exhibited the closest structural similarity with protease OmpT of *E. coli* K-12 (PDB ID: 1I78). In parallel, Foldseek identified structural homology with outer membrane protein B of *C. muridarum* strain Nigg. Supporting these findings, PANNZER annotated the protein as PorB, a well-characterized OMP (Table 2). Together, these results converge to suggest that NP_220232.1 is PorB outer membrane protein. PorB is a 38kDa protein, highly conserved across *Chlamydia* species (Kubo and Stephens 2000; Kawa and Stephens 2002) and facilitates the diffusion of dicarboxylates such as 2-oxoglutarate across the *Chlamydia* OM-crucial for metabolic activity in the context of incomplete tricarboxylic acid (TCA) cycle in *Chlamydia* (Kubo and Stephens 2001, 2000).

#### 3.1.3 Group C

**Group C** comprised four proteins: NP_219509.1 (14-stranded β-barrel), NP_219858.1 (26-stranded β-barrel), NP_219881.1 (16-stranded β-barrel), and NP_219899.1 (12-stranded β-barrel).

NP_219509.1 is annotated as ‘hypothetical protein CT_007’ in NCBI and UniProt. Structural modelling revealed a 14-stranded β-barrel protein (Figure 2). Structural alignment with DALI server revealed the best structural homology with CymA of *Klebsiella oxytoca* (PDB ID: 4V3G), as listed in Table 2. CymA, an OMP in Gram-negative bacteria, is a passive diffusion channel for large molecules (cyclodextrins and linear maltooligosaccharides) (van den Berg et al. 2015; Pajatsch et al. 1999). Complementing this, Foldseek identified a structurally similar uncharacterized protein, TC_0275, from *C. muridarum* strain Nigg, suggesting possible structural conservation. PANZZER annotated NP_219509.1 as an autotransporter beta-domain protein (Table 2). These findings support the characterization of NP_219509.1 as a β-barrel outer membrane protein, potentially involved in transport of large molecules.

NP_219858.1 is annotated as a ‘hypothetical protein CT_351’ in NCBI and UniProt. Structural modelling revealed a large β-barrel domain of 26 β-strands (Figure 2). Notably, it has a distinctive periplasmic β-jelly roll domain, similar to that observed in LptD orthologs (Figure 2). NP_219858.1 exhibited the best structural alignment with LptD (PDB ID: 4Q35) of *Shigella flexneri* (Table 2). Foldseek showed the structural alignment with an uncharacterized protein from *Chlamydiae* bacterium, while eggNOG-mapper annotated it as ‘CP_0756 (uncharacterized protein from *C. pneumoniae*), involved in the assembly of lipopolysaccharide (LPS) at the surface of outer membrane’. Additionally, annotation using PANNZER and a CDD search confirmed its classification within the LptD superfamily, further supporting its identification (Table 2 and Table S1). Combining these annotations, it can be speculated that NP_219858.1 is likely involved in LPS assembly, functioning as a LptD-like protein.

NP_219881.1 is annotated as a Porin AaxA in the UniProt database, whereas NCBI annotates it as a porin. Structural modelling revealed a 16-stranded β-barrel architecture (Figure 2). NP_219881.1 showed the highest structural homology to Porin B of *Pseudomonas putida* BIRD-1 (PDB ID: 4GEY) (Table 2). In parallel, Foldseek showed the best structural match with Porin AaxA of *C. trachomatis* L2b/UCH-1/proctitis. PANNZER and eggNOG-mapper consistently annotated the protein as Porin AaxA (Table 2). This consistent prediction across structural and sequence-based annotation tools strongly suggests that NP_219881.1 is Porin AaxA. Further, CDD search indicates that this protein’s classification within the OprB superfamily, which consists of glucose-selective porins (Saier et al. 2021) (Table S1). This finding aligns with our structure-based and sequence-based annotation prediction, i.e., porin function, likely involved in the uptake of arginine. In *C. pneumoniae*, the AaxA protein (CPn1033) functions as the outer membrane porin component of the AaxABC system-a specialized arginine-agmatine exchange complex. AaxABC system consists of three functionally linked proteins: AaxA (the outer membrane porin), AaxB (CPn1032; an arginine decarboxylase), and AaxC (CPn1031; a putative cytoplasmic membrane transporter), which work together to facilitate arginine uptake and agmatine export across cellular membrane (Kalman et al. 1999; Giles and Graham 2007). Since Chlamydial genomes lack the necessary enzymes for arginine biosynthesis, the bacterium depends on host-derived arginine for survival. Porin AaxA encodes an OM protein that facilitates the uptake of arginine into the cells and functions as a general porin with broad specificity (Smith and Graham 2008).

NP_219899.1 is annotated as DUF1207 domain-containing protein in UniProt database, while it is annotated as ‘hypothetical protein CT_389’ in NCBI. Structural modelling revealed a 12-stranded β-barrel protein with α-helical regions projecting from the extracellular side (Figure 2). DALI results showed the best structural alignment with Outer Membrane Phospholipase A (OMPLA) of *E. coli* K-12 (PDB ID: 1QD6). Foldseek exhibited the structural match with an uncharacterized protein of *Chlamydiae* bacterium. PANZZER and eggNOG-mapper classified the protein as an uncharacterized protein (Table 2). Together, these findings suggested an outer membrane-associated role, while underscoring the need for experimental annotation to elucidate its precise function.

### 3.2 Amino acid sequence variation analysis

Amino acid sequence variation analysis was performed across 106 *C. trachomatis* strains for the identified OMBB proteins. These variations were mapped onto the corresponding structural models, revealing that variations are uniformly scattered throughout the protein structural models. Notably, a significant proportion of these variations were located within the ECL region of the proteins (Table S4). Amino acid sequence variations within the ECL region suggest that these regions are under continuous selection pressure acting on the proteins at the host-pathogen interface. Variations in these regions may help the pathogen to evade immune recognition and enhance bacterial adaptability by modifying these interactions with the host environment.

### 3.3 Prediction and assessment of B-cell epitopes

ElliPro was employed to predict linear and conformational B-cell epitopes. B-cell epitopes predicted by ElliPro with a score above 0.5 were initially selected. 266 linear B-cell epitopes were then screened for homology with human proteome, and 163 epitopes with an E-value below 0.05 were excluded. The next steps involved assessing their antigenicity, allergenicity and toxicity of the remaining 103 epitopes. Further, a conservancy analysis was performed across 106 *C. trachomatis* strains. The five linear B-cell epitopes that were found to be antigenic, non-allergenic, non-toxic, non-homologous, and conserved across 106 *C. trachomatis* strains were mapped onto the structural model; however, none were located in the ECL region and thus were not considered for inclusion in the MEV design. Furthermore, 20 conformational B-cell epitopes selected in OMBB proteins based on the highest score of the tool are listed in Supplemental Table 5.

### 3.4 Prediction and assessment of T-cell epitopes

A total of 2402 CTL epitopes and 2207 HTL epitopes were predicted using the IEDB MHC I and MHC II tools, respectively, out of which 4 CTL and 29 HTL epitopes were found to be located in the ECL region (Tables-3, 4). Notably, all the selected epitopes were found to be antigenic, non-allergenic, non-toxic, non-homologous, surface-exposed, and conserved across 106 *C. trachomatis* strains (Figure 3). These T-cell epitopes were selected for inclusion in the MEV construct design.

**Figure 3:**
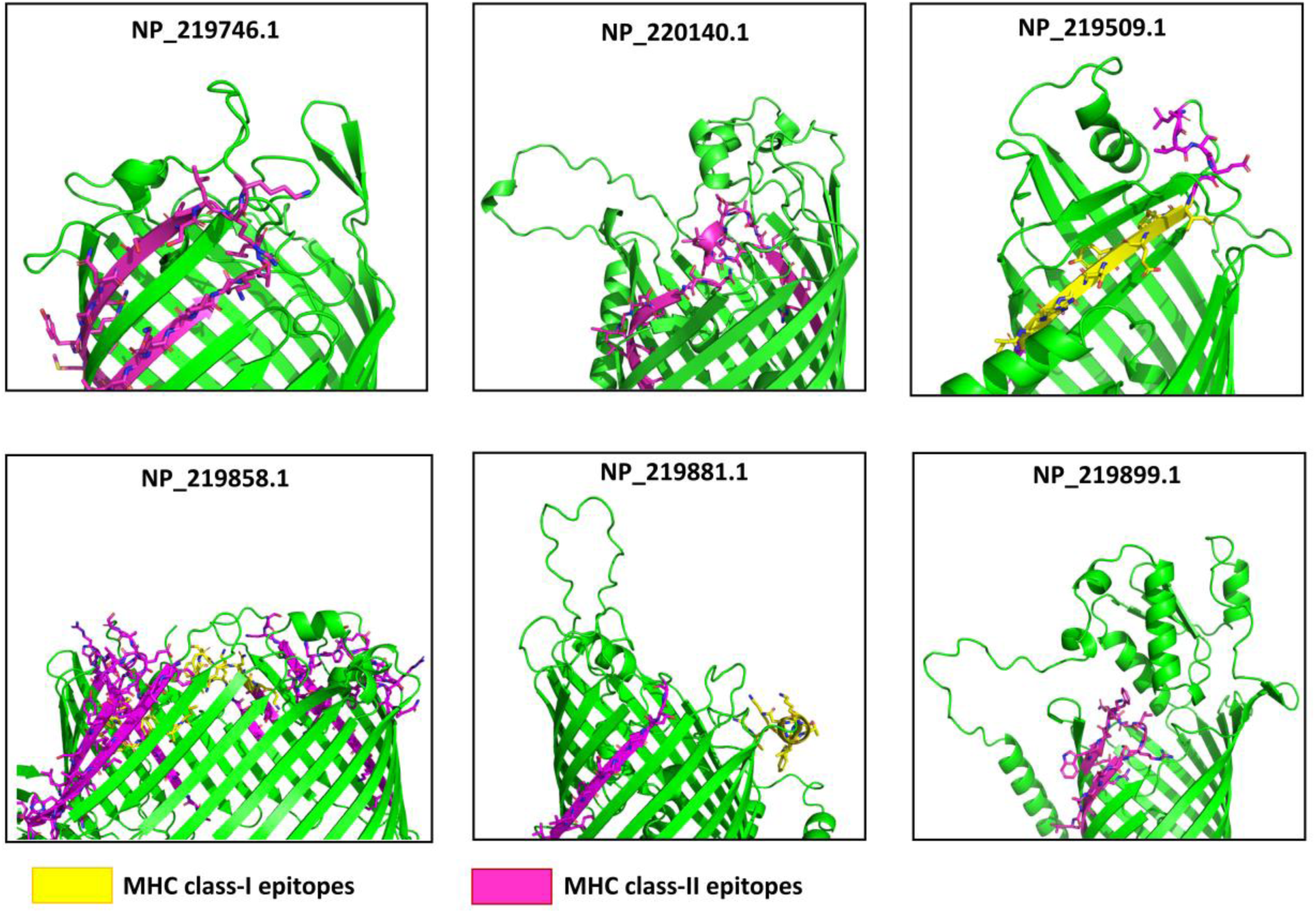
Predicted surface-exposed MHC class-I and MHC class-II epitopes. Four MHC class-I and 29 MHC class-II surface-exposed epitopes were manually identified and mapped onto structural models of the proteins: 26-stranded β-barrel protein (NP_219858.1); 16-stranded β-barrel proteins (NP_219746.1, NP_220140.1, and NP_219881.1); 14-stranded β-barrel protein (NP_219509.1); 12-stranded β-barrel protein (NP_219899.1), generated using AlphaFold 3 server.

### 3.5 Multiepitope vaccine construct design

The proposed construction of MEV is shown in Figure 4A. MEV construct design from N-terminal to C-terminal, is as follows:

EAAAKMIKLKFGVFFTVLLSSAYAHGTPQNITDLCAEYHNTQIYTLNDKIFSYTESLAGKREMAIITFKNGAIFQVEVGSQHIDSQKKAIERMKDTLRIAYLTEAKVEKLCVWNNKTPHAIAAISMANEAAAKAKFVAAWTLKAAAAAYIFEYSGREIAAYKVNSFQNVKAAYSIKWSNLTKAAYDIFPNNFSLGPGPGWILGVELDKSINKALGPGPGYGMYYRGSQTSLSLGPGPGGMYYRGSQTSLSLRKGPGPGVRYEYVEALAIPEIDGPGPGNFDGIHLYAARRISEGPGPGRYEYVEALAIPEIDGPGPGVRYEYVEALAIPEIDGPGPGNSNFDGIHLYAARRIGPGPGNFDGIHLYAARRISEGPGPGLATSEGIFEYSGREIGPGPGATSEGIFEYSGREIEGPGPGTSEGIFEYSGREIEGGPGPGSEGIFEYSGREIEGNGPGPGYLEYQMILG HKIFEHGPGPGRPVLSSDEPHIFSIKGPGPGYLEYQMILGHKIFEHGPGPGSNFSSWRFSSAHKVYGPGPGEPHIFSIKDAFHSINGPGPGHWQLFSVYEKREADKGPGPGTSSVKV NSFQNVKQEGPGPGNIKSYYAHRLAIDSSGPGPGSSVKVNSFQNVKQELGPGPGSVKVNSFQNVKQELPGPGPGSNAINIKSYYAHRLAGPGPGINIKSYYAHRLAIDSGPGPGAINIKSYYAHRLAIDGPGPGSSKLYVFGRYSGVTGGPGPGWSKFQDVGRKVRAVLGPGPGEWSKFQDVGRKVRAVKKHHHHHH

**Figure 4:**
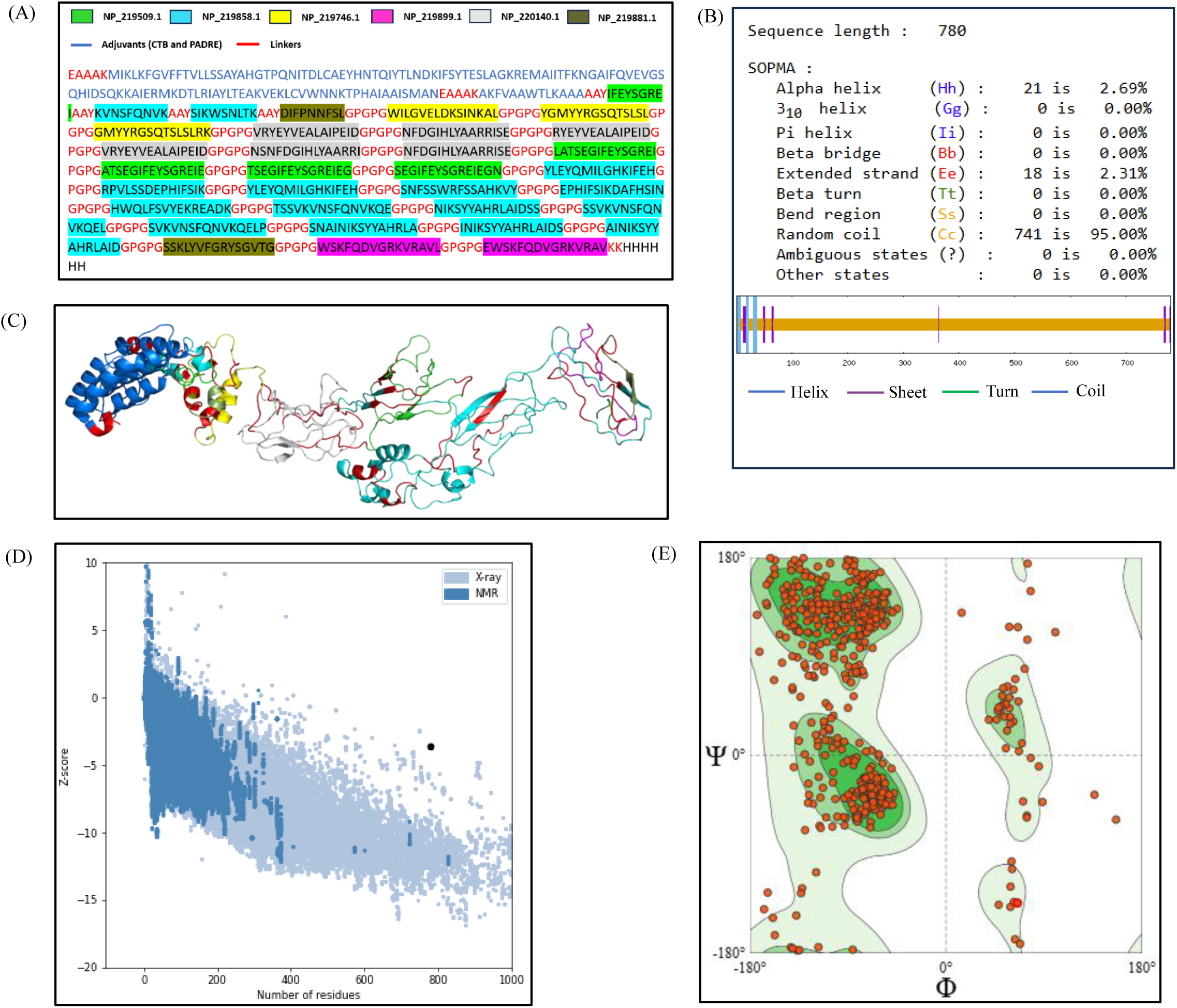
Evaluation of MEV construct. **(A)** The schematic represents the amino acid sequence of the MEV construct. **(B)** The prediction of MEV secondary structure using SOPMA tool. **(C)** The prediction of MEV tertiary structural model after refinement using GalaxyRefine server. **(D)** Z-score plot obtained from ProSA-web server. **(E)** Validation: Ramachandran plot analysis showing 84.45% of the amino acid residues falling in the favoured region.

The MEV construct consists of 780 amino acid residues.

### 3.6 Vaccine physicochemical properties and evaluation of immunological parameters

The molecular weight and isoelectric point (pI) of the vaccine were calculated to be 83.4 kDa and 8.87, respectively. In the context of vaccine design, a molecular weight below 110 kDa is generally considered suitable (Baseer et al. 2017). Similarly, the pI value of 8.87 indicates that the vaccine is slightly basic, which is acceptable, as most recombinant protein-based vaccines have pI values ranging from 5 to 9 (Vessely et al. 2009; Shone et al. 2009). The MEV comprised 780 amino acids with a molecular formula of C_3775_H_5724_N_1032_O_1096_S_10_. The instability index (II) computed through Protparam, was 29.46, classifying the MEV as a stable protein. The grand average of hydropathicity (GRAVY) score was −0.425, indicating the hydrophilic nature of the protein. In addition, MEV exhibited an antigenicity score of 0.7453 (greater than the threshold value of 0.4) and was found to be non-allergenic and non-toxic. A solubility score of 0.994, obtained through SOLpro, suggested that the protein remained soluble upon overexpression.

### 3.7 Prediction of secondary and tertiary structures

The secondary structure of MEV, predicted using SOPMA, consisted of 2.69% α-helix, 95% random coil, and 2.31% extended strand, as depicted in Figure 4B. The high proportion of random coil in the MEV indicates a higher likelihood of the protein forming antigenic epitopes (M. Yu et al. 2021). The tertiary structure was modelled using I-TASSER. The proportion of α-helix, random coil, and extended strand was consistent with the tertiary structure. This suggests that the tertiary structure predicted by I-TASSER tool is accurate. Out of five models generated by I-TASSER, we selected the one with the highest confidence and TM scores of −0.79 and 0.61±0.14, respectively. The tertiary structure of the vaccine was visualized using PyMOL, where surface-exposed HTL and CTL epitopes derived from six proteins were mapped onto the tertiary structure, as shown in Figure 4C. Further the tertiary structure was optimized using GalaxyRefine server. We selected the best-refined model based on the highest percentage of Rama-favored residues and lowest RMSD value.

### 3.8 Quality assessment of structural model

The quality of the model was assessed using ProSA-web server, which yielded a Z-score of - 3.64 for the MEV model (Figure 4D). In the Z-score plot obtained from ProSA-web server, the light blue region represents protein structures determined by X-ray crystallography, while the dark blue region corresponds to those resolved through nuclear magnetic resonance (NMR). To assess the accuracy of the predicted structure, we further evaluated the quality of the model using the Ramachandran plot tool, the structure assessment tool of SWISS-MODEL (Figure 4E). The dark green region in the Ramachandran plot represents the favored conformation, while the light green region corresponds to the allowed conformation, and the blank areas denote the disallowed regions. It was observed that 84.45% of the amino acids (i.e., dotted structures) fell within the favoured regions, confirming the structural reliability (Figure 4E). In conclusion, the predicted high-confidence tertiary structure of the MEV indicates that it is a good vaccine design.

### 3.9 Molecular docking of MEV with TLR4 immune receptor

We performed molecular docking to predict potential interactions between the vaccine construct (ligand molecule) and TLR4 receptor, based on energy minimization and structural complementarity at the receptor’s active site. Multiple docking clusters were generated, from which the top-ranked cluster was selected for analysis. The best-docked complex demonstrated a docking score of −315.45, a confidence score of 0.9647, and a ligand RMSD of 149.19 Å. The docked complex was visualized in 3D using PyMOL software (Figure 5A). The binding interactions revealed that MEV forms eleven hydrogen bonds with TLR4 within the range of 5Å interatomic distance (Figure 5B).

**Figure 5:**
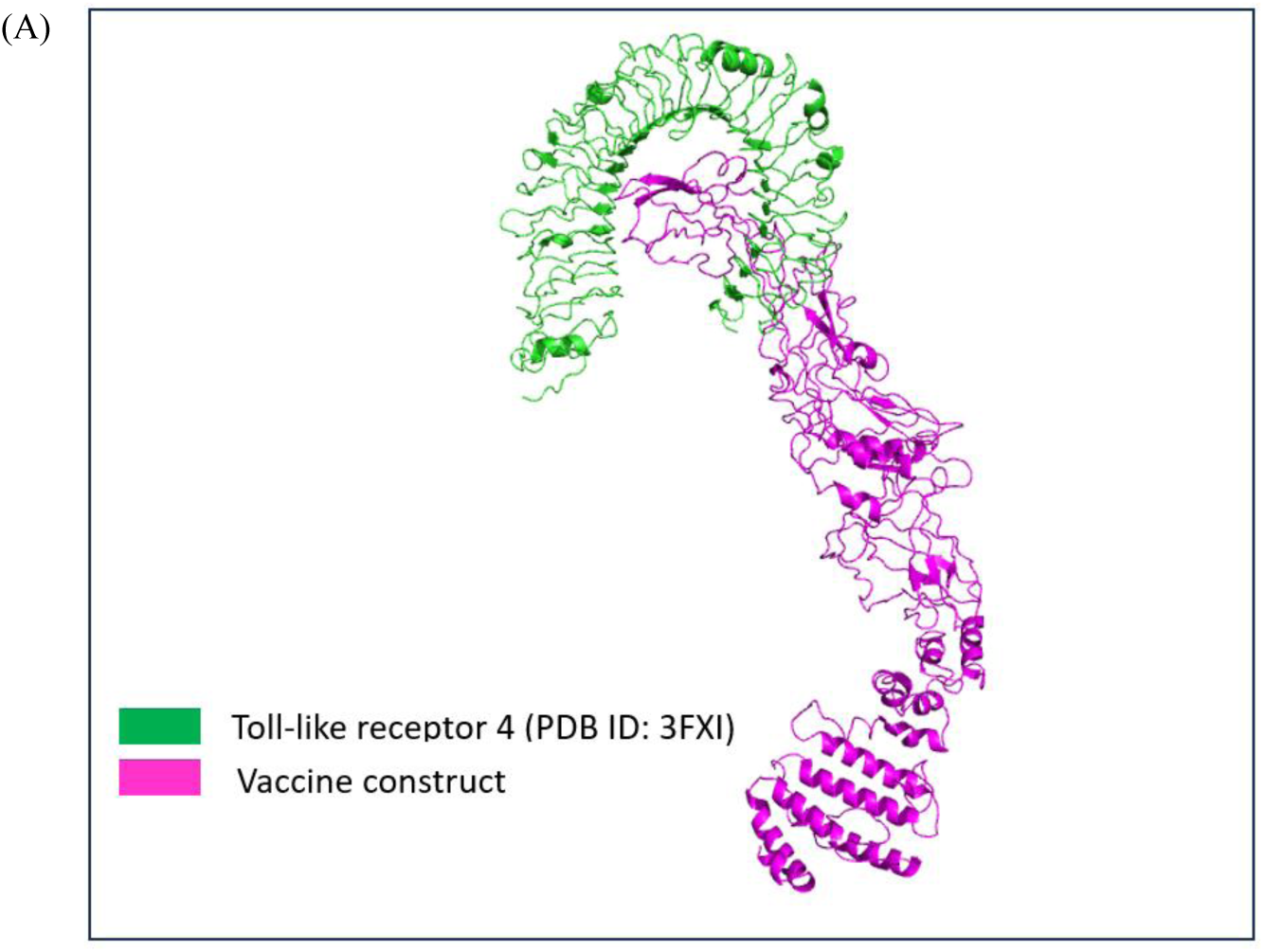
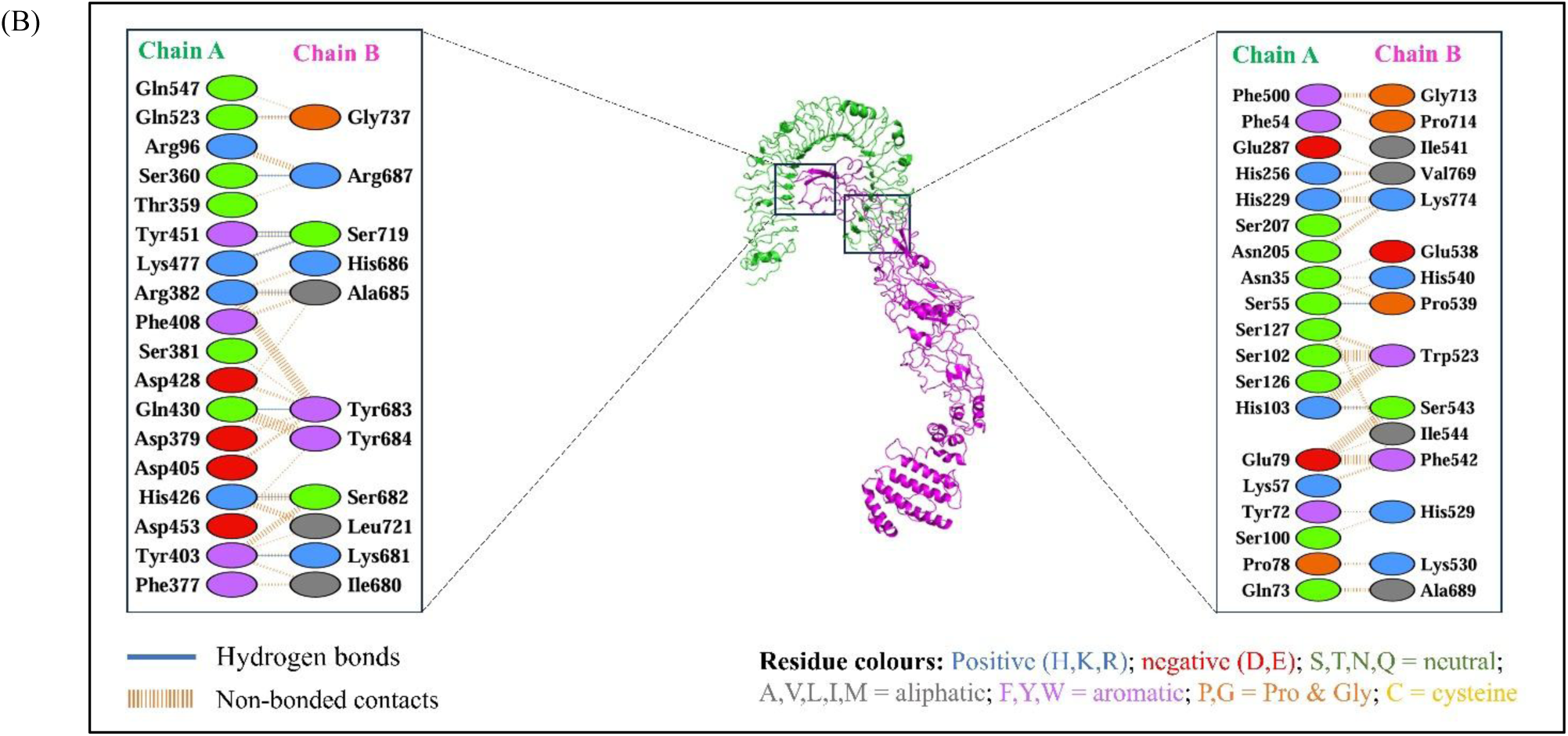
Molecular docking of MEV construct with TLR4. **(A)** The docked complex of the vaccine construct and TLR4 (PDB ID: 3FXI) is shown, obtained through HDOCK server. Since the docking of the vaccine construct was obtained with only a single chain of TLR4, we have excluded the other chain of TLR4 and myeloid differentiation factor 2 (MD-2), which is co-crystallized with the TLR4 structure. **(B)** The binding interactions between the vaccine construct and TLR4 were analysed using PDBsum server. Chain A corresponds to the TLR4 receptor, whereas Chain B corresponds to the vaccine construct. The number of hydrogen bond lines connecting two amino-acid residues of TLR4 and the vaccine construct represents the possible number of hydrogen bonds that could form between them. For non-bonded contacts, the width of the striped line corresponds to the number of atomic interactions between the residues.

## 4. Discussion

We adopted an integrated reverse vaccinology approach to systematically identify and characterize OMBB proteins of *C. trachomatis* as potential vaccine candidates. In diderm bacteria, promising vaccine targets include transmembrane β-barrel OMPs, lipoproteins anchored to the outer leaflet of the OM, and secreted proteins that interact with β-barrel OMPs (Grassmann et al. 2017). Given our focus on β-barrel OMPs, we employed a dedicated computational framework to predict and annotate them with high confidence. The prediction of antiparallel transmembrane β-sheets that form β-barrel structures is well established (Casadio et al. 2010). We relied on identifying signal peptides (SPs), β-barrel secondary structure elements, absence of transmembrane helices (TMHs), and homology to known proteins.

To ensure robust identification, nine computational tools were used to analyze the *C. trachomatis* D/UW-3/CX proteome. Specifically, four tools-OMPdb, MCMBB, TMBETADISC-RBF, and TMbed-were applied for OMBB prediction. Using consensus-based screening criteria combined with manual curation of structural models, we identified seventeen OMBB proteins. Of these, nine correspond to well-characterized Pmps described previously (PmpA to PmpI) (Cervantes et al. 2024). The remaining eight are uncharacterized OMBB proteins (NP_219746.1, NP_220140.1, NP_220200.1, NP_220232.1, NP_219509.1, NP_219858.1, NP_219881.1, and NP_219899.1). Notably, NP_220200.1 encodes the extensively studied MOMP protein, serving as an internal validation of our computational pipeline. For structural modelling, we employed five independent prediction tools-AlphaFold 3, ESMFold, SWISS-MODEL, RoseTTAFold, and TrRosetta. The resulting models showed strong concordance, with RMSD values under 5 Å when aligned using the US-Align server, indicating minimal structural deviation and high model accuracy. While these in silico predictions are promising, experimental validation via techniques such as X-ray crystallography or NMR spectroscopy remains essential. Our structure-based functional annotation approach leverages the principle that protein structures are more conserved than sequences over evolutionary timescales (Chothia and Lesk 1986), allowing functional inference even when sequence similarity is low. By integrating both structure- and sequence-based annotation methods, we achieved more robust predictions, with several OMBB candidates showing homology to known OMPs such as BamA, OmpA, PorB, and LptD. Furthermore, analysis of amino acid sequence variation within the ECL regions revealed evidence of ongoing selection pressure, likely driven by immune evasion strategies at the host– pathogen interface. These variations may facilitate immune recognition escape and promote bacterial adaptability, underscoring the importance of targeting ECL regions in vaccine design.

Recognizing that these identified OMPs are surface-exposed and immunogenic, we evaluated their potential as vaccine candidates through epitope prediction and MEV design. HTL and CTL epitopes were linked using flexible GPGPG and AAY linkers, respectively, which help maintain proper folding and domain separation (X. Chen, Zaro, and Shen 2013). Flexible linkers such as GPGPG, AAY, and KK contain residues like Gly and Ser to promote structural flexibility and favorable interdomain interactions, whereas rigid linkers such as EAAAK maintain spacing and reduce undesirable interactions. Adjuvants are essential for enhancing the immunogenicity of MEVs. In this study, CTB and PADRE peptide were incorporated as adjuvants. CTB has been widely used for its ability to boost vaccine responses (Hou et al. 2014), while PADRE serves as a universal T-helper epitope capable of inducing robust, broad-spectrum MHC-II responses and Th1 polarization (Ghaffari-Nazari et al. 2015). The inclusion of these adjuvants enhances the stability, potency, and durability of the vaccine construct (Lee and Nguyen 2015).

For early-stage vaccine design, molecular weight is a critical parameter; proteins under 110 kDa are generally favoured (Baseer et al. 2017). Our MEV construct comprises 780 amino acids with a molecular weight of 83.4 kDa, well within this recommended range. Solubility assessment predicted high solubility in *E. coli* (probability 0.994), supporting feasibility for recombinant production (Khatoon, Pandey, and Prajapati 2017). Antigenicity (0.7453) exceeded threshold values, and allergenicity predictions indicated no allergenic potential. Physicochemical properties, including aliphatic index, GRAVY score, and instability index, suggested thermostability, hydrophilicity, and overall protein stability. Secondary structure prediction revealed ∼95% random coils, a feature associated with enhanced epitope presentation potential (M. Yu et al. 2021). Tertiary structure modelling using I-TASSER yielded an estimated TM-score of 0.61 ± 0.14, indicating a correctly predicted topology (TM-score >0.5) (Y. Zhang 2008). The C-score of −0.79 fell well within acceptable limits (−5 to 2), further supporting model reliability. Structural validation using ProSA-web provided a Z-score consistent with experimentally determined structures, while Ramachandran plot analysis showed 84.45% of residues in favoured regions. Beyond static structure, vaccine–immune receptor interactions are critical. Toll-like receptor 4 (TLR4) recognizes pathogen-associated molecular patterns such as LPS from *C. trachomatis* and is central to innate immune responses (Mei, Li, and Zuo 2021; Joyee and Yang 2008). Molecular docking analyses revealed stable interactions at the atomic interface between TLR4 and the MEV construct, suggesting the vaccine is capable of effective immune receptor engagement. In summary, this study presents a comprehensive computational pipeline for the identification of β-barrel OMPs and the design of a MEV construct. The approach combines sequence- and structure-based analyses, epitope prediction, rational linker design, adjuvant inclusion, and structural validation to develop a promising vaccine candidate. While the in silico results strongly support its potential, experimental validation is essential to confirm its safety, immunogenicity, and protective efficacy.

## 5. Conclusion

In this study, we employed an integrated computational pipeline to identify OMBB proteins of *C. trachomatis* as promising vaccine candidates and designed an MEV incorporating epitopes from six OMBB proteins. The MEV construct demonstrated favourable physicochemical properties, predicted solubility, antigenicity, and structural stability, supporting its potential as a vaccine candidate. However, while these in silico results are encouraging, further experimental validation through in vitro and in vivo studies, along with subsequent pharmacological evaluation, will be essential to confirm its safety, immunogenicity, and protective efficacy.

## Acknowledgements

AP is a recipient of junior research fellowship from the Department of Biotechnology, Government of India. JK is a recipient of junior research fellowship from the University Grants Commission, Government of India.

## Author contributions statement

**Amisha Panda:** methodology, software, investigation, data analysis, data curation, writing— original draft preparation. **Jahnvi Kapoor:** data analysis, visualization, writing—reviewing and editing. **B. Hareramadas:** visualization, writing—reviewing and editing. **Ilmas Naqvi:** visualization, writing—reviewing and editing. **Satish Ganta:** visualization, writing— reviewing and editing. **Ravindresh Chhabra:** data analysis, writing—original draft preparation, writing—reviewing and editing. **Sanjiv Kumar:** conceptualization, methodology, software, writing—reviewing and editing, supervision. **Anannya Bandyopadhyay:** methodology, data analysis, writing—original draft preparation, writing—reviewing and editing, supervision. All authors reviewed the final draft of the manuscript. All authors contributed to the manuscript revision, read and approved the submitted version.

## Data Availability Statement

The data supporting the findings of this study are available from the corresponding author upon reasonable request.

