## Supplemental material for "Design of a Multi-Epitope Vaccine using β-barrel Outer Membrane Proteins Identified in *Chlamydia trachomatis*"

**Supplemental Tables**

**Table S1**: Comprehensive results from computational tools employed in our study to predict outer membrane β-barrel proteins.

| **Sequence Identifier** | **Protein name≠** | **Pepstats** | | **SPAAN** | **CDD Search** | **SignalP** | | **PSORTb** | | **CELLO** | **OMPdb** | **TMBETADISC-RBF** | **MCMBB** | **TMbed** | |
| --- | --- | --- | --- | --- | --- | --- | --- | --- | --- | --- | --- | --- | --- | --- | --- |
|  |  | **AA** | **Mol Wt. (kDa)** |  |  | **Signal Peptide** | **CS Position** | **Localization** | **Score** |  |  |  |  | **TMbedcount_b** | **TMbedcount_B** |
| **GROUP A** |  |  |  |  |  |  |  |  |  |  |  |  |  |  |  |
| NP_219746.1 | Outer membrane protein assembly factor BamA | 792 | 88.8 | 0.50 | BamA | SP(Sec/SPI) | CS pos: 26-27. | Outer Membrane | 10 | Outer Membrane | 524 | Outer Membrane Protein | 0.004 | 68 | 68 |
| NP_219923.1 | Outer membrane protein PmpB | 1751 | 183.0 | 0.91 | ChlamPMP_M | SP(Sec/SPI) | CS pos: 19-20. | Outer Membrane | 9.99 | Extracellular | 14 | Outer Membrane Protein | 0.024 | 57 | 52 |
| NP_219924.1 | Outer membrane protein PmpC | 1770 | 187.0 | 0.85 | ChlamPMP_M | SP(Sec/SPI) | CS pos: 20-21. | Outer Membrane | 9.99 | Extracellular | 17 | Outer Membrane Protein | 0.018 | 56 | 55 |
| NP_220392.1 | Outer membrane protein PmpF | 1034 | 112.4 | 0.85 | ChlamPMP_M | SP(Sec/SPI) | CS pos: 25-26 | Extracellular | 10 | Outer Membrane | 7 | Outer Membrane Protein | 0.014 | 56 | 54 |
| NP_220393.1 | Outer membrane protein PmpG | 1013 | 107.3 | 0.90 | ChlamPMP_M | SP(Sec/SPI) | CS pos: 27-28 | Outer Membrane | 9.99 | Outer Membrane | 15 | Outer Membrane Protein | 0.016 | 55 | 53 |
| NP_220394.1 | Outer membrane protein PmpH | 1016 | 107.9 | 0.87 | ChlamPMP_M | SP(Sec/SPI) | CS pos: 24-25 | Extracellular | 9.46 | Extracellular | 12 | Outer Membrane Protein | 0/011 | 57 | 54 |
| **GROUP B** |  |  |  |  |  |  |  |  |  |  |  |  |  |  |  |
| NP_219922.1 | Outer membrane protein PmpA | 975 | 105.6 | 0.80 | ChlamPMP_M | OTHER | 0 | Unknown | 7 | Extracellular | 14 | Outer Membrane Protein | -0.011 | 56 | 54 |
| NP_220140.1 | CHLPN 76kDa Homolog | 432 | 48.3 | 0.62 | - | SP(Sec/SPI) | CS pos: 25-26. | Unknown | 2.5 | Outer Membrane | 0 | Outer Membrane Protein | 0.007 | 73 | 68 |
| NP_220200.1 | Major outer membrane porin, serovar D | 393 | 42.4 | 0.78 | Chlam_OMP | SP(Sec/SPI) | CS pos: 22-23. | Outer Membrane | 10 | Outer Membrane | 500 | Outer Membrane Protein | -0.014 | 45 | 43 |
| NP_220232.1 | Outer membrane protein B | 340 | 37.4 | 0.72 | Chlam_OMP superfamily | SP(Sec/SPI) | CS pos: 26-27. | Outer Membrane | 9.93 | Outer Membrane | 368 | Outer Membrane Protein | -0.012 | 39 | 36 |
| NP_220332.1 | Outer membrane protein PmpD | 1531 | 160.7 | 0.75 | ChlamPMP_M | LIPO(Sec/SPII) | CS pos: 31-32. | Outer Membrane | 9.99 | Outer Membrane | 17 | Outer Membrane Protein | -0.009 | 62 | 57 |
| NP_220391.1 | Outer membrane protein PmpE | 964 | 104.7 | 0.90 | ChlamPMP_M | SP(Sec/SPI) | CS pos: 18-19. | Outer Membrane | 9.83 | Extracellular | 13 | Outer Membrane Protein | -0.003 | 57 | 54 |
| NP_220396.1 | Outer membrane protein PmpI | 878 | 95.6 | 0.68 | Autotransporter | SP(Sec/SPI) | CS pos: 24-25. | Extracellular | 9.43 | Outer Membrane | 12 | Outer Membrane Protein | -0.017 | 55 | 52 |
| **GROUP C** |  |  |  |  |  |  |  |  |  |  |  |  |  |  |  |
| NP_219509.1 | Uncharacterized protein CT_007 | 316 | 35.7 | 0.45 | - | LIPO(Sec/SPII) | CS pos: 26-27. | Unknown | 2 | Outer Membrane | 0 | Outer Membrane Protein | -0.006 | 69 | 54 |
| NP_219858.1 | Uncharacterized protein | 697 | 80.0 | 0.30 | LptD superfamily | SP(Sec/SPI) | CS pos: 21-22. | Unknown | 2 | Outer Membrane | 33 | Non-Outer Membrane Protein | -0.051 | 103 | 101 |
| NP_219881.1 | Porin AaxA | 442 | 49.4 | 0.60 | OprB superfamily | OTHER | 0 | Extracellular | 9.64 | Outer Membrane | 36 | Non-Outer Membrane Protein | -0.004 | 69 | 62 |
| NP_219899.1 | DUF1207 domain-containing protein | 408 | 47.0 | 0.13 | DUF1207 superfamily | SP(Sec/SPI) | CS pos: 24-25. | Cytoplasmic | 8.96 | Cytoplasmic | 1 | Non-Outer Membrane Protein | -0.081 | 56 | 50 |

^≠^ Proteins were named as per their annotation in NCBI and UniProt databases, searched using Sequence Identifier.

**TableS2:** Structural alignment of structural models of the selected OMBB proteins, generated by five modelling tools

| **Sequence Identifier** | **Protein name^≠^** | **Confidence Scores** | | | | | **Structural alignment**  **(US-Align)** | |
| --- | --- | --- | --- | --- | --- | --- | --- | --- |
|  |  | **AlphaFold 3** | **RoseTTAFold** | **TrRosetta** | **ESMFold** | **SWISS-MODEL** | **Figure** | **RMSD**  **(Å)** |
| **GROUP A** |  |  |  |  |  |  |  |  |
| NP_219746.1^*^ | Outer membrane protein assembly factor BamA | 0.71 | 0.80 | 0.827 | 0.788 | 0.89 | 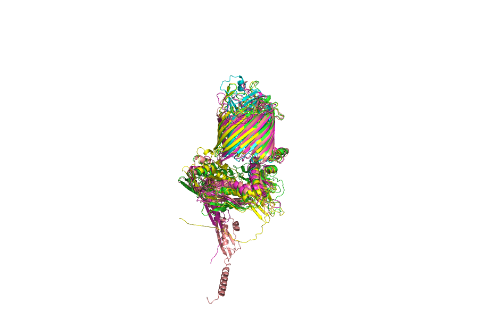 | 4.92 |
| **GROUP B** |  |  |  |  |  |  |  |  |
| NP_220140.1 | CHLPN 76kDa Homolog | 0.82 | 0.64 | 0.812 | 0.785 | 0.77 | 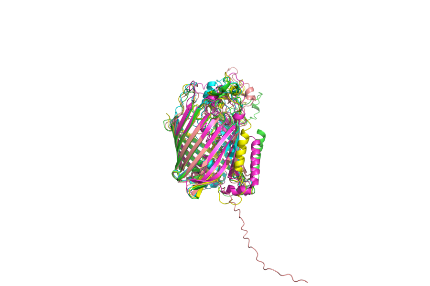 | 3.55 |
| NP_220200.1 | Major outer membrane porin, serovar D | 0.85 | 0.59 | 0.739 | 0.623 | 0.77 | 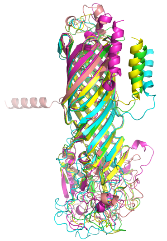 | 3.82 |
| NP_220232.1 | Outer membrane protein B | 0.78 | 0.66 | 0.832 | 0.655 | 0.69 | 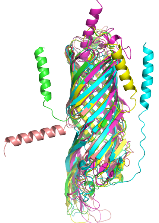 | 3.46 |
| **GROUP C** |  |  |  |  |  |  |  |  |
| NP_219509.1 | Hypothetical protein CT_007 | 0.83 | 0.71 | 0.875 | 0.833 | 0.85 | 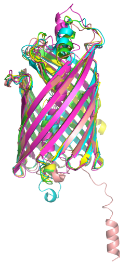 | 2.10 |
| NP_219858.1 | Hypothetical protein CT_351 | 0.84 | 0.72 | 0.818 | 0.655 | 0.84 | 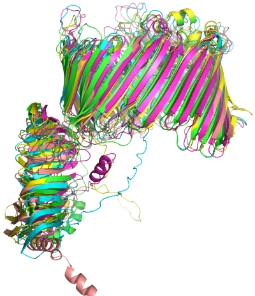 | 4.00 |
| NP_219881.1 | Porin AaxA | 0.89 | 0.66 | 0.761 | 0.746 | 0.82 | 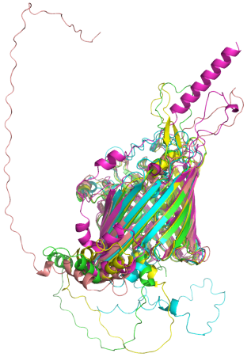 | 2.95 |
| NP_219899.1 | DUF1207 domain-containing protein | 0.88 | 0.69 | 0.740 | 0.710 | 0.84 | 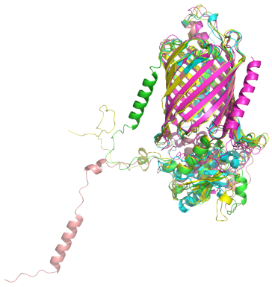 | 3.33 |

**^≠^** Proteins were named as per their annotation in NCBI and UniProt databases, searched using Sequence Identifier.

^*^Due to an error in RoseTTAFold when submitting the full FASTA sequence, the entire structural model could not be generated. Therefore, a structural model was created using only 500 amino acid residues.

Green color, cyan color, magenta color, yellow color and salmon color represents the structural models generated by AlphaFold 3, ESMFold, RoseTTAFold, SWISS-MODEL and TrRosetta, respectively.

**TableS3:** *C. trachomatis* 106 strains searched for predicted OMP’s variations compared to the reference genome D/UW-3/CX.

| **Assembly Accession** | **Assembly Name** | **Organism Name** | **Strain Name** | **Genome Size (Mb)** | **Scaffolds** | **CDS^#^** |
| --- | --- | --- | --- | --- | --- | --- |
| GCF_000008725.1 | ASM872v1 | *Chlamydia trachomatis* | D/UW-3/CX | 1.042519 | 1 | 887 |
| GCF_000012125.1 | ASM1212v1 | *Chlamydia trachomatis* | A/HAR-13 | 1.051969 | 2 | 888 |
| GCF_000026905.1 | ASM2690v1 | *Chlamydia trachomatis* | B/TZ1A828/OT | 1.044282 | 1 | 883 |
| GCF_000026925.1 | ASM2692v1 | *Chlamydia trachomatis* | Jali20 | 1.044352 | 1 | 884 |
| GCF_000068525.2 | ASM6852v2 | *Chlamydia trachomatis* | UCH-1 | 1.038863 | 1 | 880 |
| GCF_000068585.1 | ASM6858v1 | *Chlamydia trachomatis* | L2/434/Bu | 1.038842 | 1 | 879 |
| GCF_000092485.1 | ASM9248v1 | *Chlamydia trachomatis* | D-EC | 1.050015 | 2 | 909 |
| GCF_000092665.1 | ASM9266v1 | *Chlamydia trachomatis* | E/150 | 1.042996 | 1 | 890 |
| GCF_000092685.1 | ASM9268v1 | *Chlamydia trachomatis* | G/9768 | 1.04281 | 1 | 895 |
| GCF_000092705.1 | ASM9270v1 | *Chlamydia trachomatis* | G/11222 | 1.042354 | 1 | 898 |
| GCF_000092725.1 | ASM9272v1 | *Chlamydia trachomatis* | G/11074 | 1.042875 | 1 | 897 |
| GCF_000092745.1 | ASM9274v1 | *Chlamydia trachomatis* | E/11023 | 1.043025 | 1 | 893 |
| GCF_000092805.1 | ASM9280v1 | *Chlamydia trachomatis* | G/9301 | 1.042811 | 1 | 895 |
| GCF_000093005.1 | ASM9300v1 | *Chlamydia trachomatis* | D-LC | 1.050013 | 2 | 908 |
| GCF_000173475.1 | ASM17347v1 | *Chlamydia trachomatis* | 6276 | 1.043181 | 1 | 897 |
| GCF_000173495.1 | ASM17349v1 | *Chlamydia trachomatis* | 6276s | 1.043182 | 1 | 896 |
| GCF_000173515.1 | ASM17351v1 | *Chlamydia trachomatis* | 70 | 1.048006 | 1 | 885 |
| GCF_000173535.1 | ASM17353v1 | *Chlamydia trachomatis* | 70s | 1.046064 | 1 | 880 |
| GCF_000174055.1 | ASM17405v1 | *Chlamydia trachomatis* | 2923 | 1.042757 | 1 | 890 |
| GCF_000175515.1 | ASM17551v1 | *Chlamydia trachomatis* | L2tet1 | 1.076624 | 1 | 908 |
| GCF_000210495.1 | ASM21049v1 | *Chlamydia trachomatis* | Sweden2 | 1.042839 | 1 | 891 |
| GCF_000220105.1 | ASM22010v1 | *Chlamydia trachomatis* | L2c | 1.038313 | 1 | 879 |
| GCF_000226605.1 | ASM22660v1 | *Chlamydia trachomatis* | A2497 | 1.051806 | 2 | 890 |
| GCF_000284475.1 | ASM28447v1 | *Chlamydia trachomatis* | A2497 | 1.044325 | 1 | 881 |
| GCF_000304495.1 | Cm_ESW3 | *Chlamydia trachomatis* | E/SW3 | 1.042903 | 1 | 892 |
| GCF_000304515.1 | Cm_FSW4 | *Chlamydia trachomatis* | F/SW4 | 1.042736 | 1 | 892 |
| GCF_000304535.1 | Cm_FSW5 | *Chlamydia trachomatis* | F/SW5 | 1.042743 | 1 | 894 |
| GCF_000318525.1 | ASM31852v2 | *Chlamydia trachomatis* | A/363 | 1.051528 | 2 | 890 |
| GCF_000318565.1 | ASM31856v1 | *Chlamydia trachomatis* | A/7249 | 1.051879 | 2 | 889 |
| GCF_000318645.1 | ASM31864v1 | *Chlamydia trachomatis* | E/Bour | 1.050091 | 2 | 898 |
| GCF_000318765.1 | ASM31876v1 | *Chlamydia trachomatis* | Ia/SotonIa1 | 1.05013 | 2 | 903 |
| GCF_000318785.1 | ASM31878v1 | *Chlamydia trachomatis* | Ia/SotonIa3 | 1.050283 | 2 | 903 |
| GCF_000318825.1 | ASM31882v1 | *Chlamydia trachomatis* | L1/440/LN | 1.039261 | 1 | 879 |
| GCF_000318845.1 | ASM31884v1 | *Chlamydia trachomatis* | L1/115 | 1.046584 | 2 | 887 |
| GCF_000318865.1 | ASM31886v1 | *Chlamydia trachomatis* | L1/224 | 1.046319 | 2 | 888 |
| GCF_000318885.1 | ASM31888v1 | *Chlamydia trachomatis* | L2/25667R | 1.038839 | 1 | 880 |
| GCF_000318905.1 | ASM31890v1 | *Chlamydia trachomatis* | L2b/8200/07 | 1.046364 | 2 | 889 |
| GCF_000318925.1 | ASM31892v1 | *Chlamydia trachomatis* | L2b/UCH-2 | 1.046337 | 2 | 889 |
| GCF_000318985.1 | ASM31898v1 | *Chlamydia trachomatis* | L2b/795 | 1.04622 | 2 | 889 |
| GCF_000319105.1 | ASM31910v1 | *Chlamydia trachomatis* | L3/404/LN | 1.046672 | 2 | 886 |
| GCF_000364765.1 | ASM36476v1 | *Chlamydia trachomatis* | L2/434/Bu(i) | 1.046341 | 2 | 886 |
| GCF_000364785.1 | ASM36478v1 | *Chlamydia trachomatis* | L2/434/Bu(f) | 1.046341 | 2 | 886 |
| GCF_000441585.1 | ASM44158v1 | *Chlamydia trachomatis* | RC-F/69 | 1.046087 | 1 | 882 |
| GCF_000441615.1 | ASM44161v1 | *Chlamydia trachomatis* | RC-L2(s)/46 | 1.038817 | 1 | 879 |
| GCF_000441635.1 | ASM44163v1 | *Chlamydia trachomatis* | RC-F(s)/852 | 1.083893 | 1 | 907 |
| GCF_000441655.1 | ASM44165v1 | *Chlamydia trachomatis* | RC-J/943 | 1.038797 | 1 | 880 |
| GCF_000441675.1 | ASM44167v1 | *Chlamydia trachomatis* | RC-J/953 | 1.038792 | 1 | 884 |
| GCF_000441695.1 | ASM44169v1 | *Chlamydia trachomatis* | RC-L2(s)/3 | 1.046171 | 1 | 880 |
| GCF_000441715.1 | ASM44171v1 | *Chlamydia trachomatis* | RC-F(s)/342 | 1.083842 | 1 | 908 |
| GCF_000441735.1 | ASM44173v1 | *Chlamydia trachomatis* | RC-J(s)/122 | 1.061854 | 1 | 901 |
| GCF_000441755.1 | ASM44175v1 | *Chlamydia trachomatis* | RC-J/966 | 1.038839 | 1 | 881 |
| GCF_000441775.1 | ASM44177v1 | *Chlamydia trachomatis* | J/6276tet1 | 1.080953 | 1 | 923 |
| GCF_000441795.1 | ASM44179v1 | *Chlamydia trachomatis* | RC-J/971 | 1.038442 | 1 | 881 |
| GCF_000441815.1 | ASM44181v1 | *Chlamydia trachomatis* | RC-L2/55 | 1.03887 | 1 | 880 |
| GCF_000507225.1 | ASM50722v1 | *Chlamydia trachomatis* | C/TW-3 | 1.051055 | 2 | 888 |
| GCF_000590575.1 | ASM59057v1 | *Chlamydia trachomatis* | F/1-93 | 1.042588 | 1 | 885 |
| GCF_000590615.1 | ASM59061v1 | *Chlamydia trachomatis* | F/6-94 | 1.042767 | 1 | 893 |
| GCF_000590635.1 | ASM59063v1 | *Chlamydia trachomatis* | F/11-96 | 1.042853 | 1 | 890 |
| GCF_000590675.1 | ASM59067v1 | *Chlamydia trachomatis* | D/13-96 | 1.042752 | 1 | 894 |
| GCF_000590755.1 | ASM59075v1 | *Chlamydia trachomatis* | J/31-98 | 1.043207 | 1 | 896 |
| GCF_000971705.1 | ASM97170v1 | *Chlamydia trachomatis* | L2b/CS19/08 | 1.046364 | 2 | 888 |
| GCF_000971725.1 | ASM97172v1 | *Chlamydia trachomatis* | L2b/CS784/08 | 1.046364 | 2 | 888 |
| GCF_001183765.1 | ASM118376v1 | *Chlamydia trachomatis* | D/CS637/11 | 1.050173 | 2 | 907 |
| GCF_001183805.1 | ASM118380v1 | *Chlamydia trachomatis* | E/CS1025/11 | 1.050536 | 2 | 900 |
| GCF_001183825.1 | ASM118382v1 | *Chlamydia trachomatis* | F/CS847/08 | 1.050553 | 2 | 900 |
| GCF_001183845.1 | ASM118384v1 | *Chlamydia trachomatis* | Ia/CS190/96 | 1.049505 | 2 | 910 |
| GCF_001655335.1 | ASM165533v1 | *Chlamydia trachomatis* | E-103 | 1.050521 | 2 | 900 |
| GCF_001655375.1 | ASM165537v1 | *Chlamydia trachomatis* | E-160 | 1.050509 | 2 | 899 |
| GCF_001655415.1 | ASM165541v1 | *Chlamydia trachomatis* | E-547 | 1.050505 | 2 | 898 |
| GCF_001655455.1 | ASM165545v1 | *Chlamydia trachomatis* | E-8873 | 1.05021 | 2 | 901 |
| GCF_001655495.1 | ASM165549v1 | *Chlamydia trachomatis* | E-32931 | 1.056419 | 2 | 899 |
| GCF_001655535.1 | ASM165553v1 | *Chlamydia trachomatis* | E-DK-20 | 1.055535 | 2 | 899 |
| GCF_001655575.1 | ASM165557v1 | *Chlamydia trachomatis* | F-6068 | 1.050231 | 2 | 902 |
| GCF_001885175.1 | ASM188517v1 | *Chlamydia trachomatis* | QH111L | 1.033342 | 2 | 882 |
| GCF_002088315.1 | ASM208831v1 | *Chlamydia trachomatis* | chxRP4F1 | 1.042393 | 1 | 897 |
| GCF_002088335.1 | ASM208833v1 | *Chlamydia trachomatis* | chxRP4D10 | 1.042369 | 1 | 903 |
| GCF_002088355.1 | ASM208835v1 | *Chlamydia trachomatis* | chxRP1H1 | 1.042376 | 1 | 896 |
| GCF_002776745.1 | ASM277674v1 | *Chlamydia trachomatis* | SQ32 | 1.047716 | 1 | 889 |
| GCF_002776795.1 | ASM277679v1 | *Chlamydia trachomatis* | SQ29 | 1.047716 | 1 | 889 |
| GCF_002776845.1 | ASM277684v1 | *Chlamydia trachomatis* | SQ20 | 1.04853 | 1 | 902 |
| GCF_002776885.1 | ASM277688v1 | *Chlamydia trachomatis* | SQ24 | 1.048538 | 1 | 902 |
| GCF_002776955.1 | ASM277695v1 | *Chlamydia trachomatis* | SQ09 | 1.042567 | 1 | 884 |
| GCF_002776975.1 | ASM277697v1 | *Chlamydia trachomatis* | SQ10 | 1.042575 | 1 | 887 |
| GCF_002776995.1 | ASM277699v1 | *Chlamydia trachomatis* | SQ12 | 1.042577 | 1 | 885 |
| GCF_002777015.1 | ASM277701v1 | *Chlamydia trachomatis* | SQ14 | 1.042578 | 1 | 885 |
| GCF_002777035.1 | ASM277703v1 | *Chlamydia trachomatis* | SQ01 | 1.048696 | 1 | 897 |
| GCF_002777055.1 | ASM277705v1 | *Chlamydia trachomatis* | SQ02 | 1.048696 | 1 | 897 |
| GCF_002777075.1 | ASM277707v1 | *Chlamydia trachomatis* | SQ05 | 1.048696 | 1 | 897 |
| GCF_002777095.1 | ASM277709v1 | *Chlamydia trachomatis* | SQ25 | 1.043246 | 1 | 891 |
| GCF_002777135.1 | ASM277713v1 | *Chlamydia trachomatis* | SQ15 | 1.048307 | 1 | 898 |
| GCF_002777155.1 | ASM277715v1 | *Chlamydia trachomatis* | SQ19 | 1.048306 | 1 | 898 |
| GCF_004135145.1 | ASM413514v1 | *Chlamydia trachomatis* | tet9 | 1.054567 | 1 | 899 |
| GCF_018336255.1 | ASM1833625v1 | *Chlamydia trachomatis* | L2/Hefty/ct350::Tn | 1.040265 | 1 | 880 |
| GCF_018336275.1 | ASM1833627v1 | *Chlamydia trachomatis* | L2/Hefty/ct732::Tn | 1.040265 | 1 | 881 |
| GCF_018336295.1 | ASM1833629v1 | *Chlamydia trachomatis* | L2/Hefty/ct054::Tn | 1.040265 | 1 | 881 |
| GCF_018336315.1 | ASM1833631v1 | *Chlamydia trachomatis* | L2/Hefty/ct404::Tn | 1.040265 | 1 | 880 |
| GCF_018337015.1 | ASM1833701v1 | *Chlamydia trachomatis* | L2/Hefty/ct590::Tn | 1.040265 | 1 | 880 |
| GCF_025723025.1 | ASM2572302v1 | *Chlamydia trachomatis* | L2f | 1.045668 | 2 | 887 |
| GCF_036285905.1 | ASM3628590v1 | *Chlamydia trachomatis* | LGV II 434 | 1.039264 | 1 | 879 |
| GCF_039778305.1 | ASM3977830v1 | *Chlamydia trachomatis* | BQ05_F_6370 | 1.048006 | 1 | 886 |
| GCF_039778315.1 | ASM3977831v1 | *Chlamydia trachomatis* | BQ05_M04431_F_6370 | 1.048006 | 1 | 886 |
| GCF_039778325.1 | ASM3977832v1 | *Chlamydia trachomatis* | BQ06_D0_J_p225 | 1.0427 | 1 | 899 |
| GCF_039778335.1 | ASM3977833v1 | *Chlamydia trachomatis* | BQ06_K06160_J_p225 | 1.042701 | 1 | 898 |
| GCF_039778345.1 | ASM3977834v1 | *Chlamydia trachomatis* | BQ06_L04117_J_p225 | 1.042701 | 1 | 898 |
| GCF_039778415.1 | ASM3977841v1 | *Chlamydia trachomatis* | BQ07_A11257_D_6319 | 1.042473 | 1 | 894 |
| GCF_039778425.1 | ASM3977842v1 | *Chlamydia trachomatis* | BQ07_D0_D_6319 | 1.042473 | 1 | 894 |

^#^CDS: Coding Sequences

**TableS4:** Amino acid sequence variations identified in the different regions of the selected eight transmembrane beta-barrel proteins.

| **Sequence Identifier** | **Protein name^≠^** | **Total number** | **Amino acid sequence variations** | **Variations in ECL region** | **Variations in ICL region** | **Variations in TM region** | **Variations in other region** |
| --- | --- | --- | --- | --- | --- | --- | --- |
| **GROUP A** |  |  |  |  |  |  |  |
| NP_219746.1 | Outer membrane protein assembly factor BamA | 16 | S21, R22, D79, A94, Y129, V343, P355, G393, R413, P575, S631, G657, T693, Y734, A745, P769 | P769, T693, P575, G657, Y734 | A745, S631, | None | S21, R22, D79, A94, Y129, V343, P355, G393, R413 |
| **GROUP B** |  |  |  |  |  |  |  |
| NP_220140.1 | CHLPN 76kDa Homolog | 11 | G8, V58, L76, A83, M86, T89, V229, A246, T306, T336, A366 | A83, M86, T89, A246, T306, A366 | None | L76, V229, T336, V58 | G8 |
| NP_220200.1^∞^ | Major outer membrane porin, serovar D | 93 | A52, V62, K75, Q83, A86, K87, P88, T89, T90, D91, T92, G93, N94, S95, A96, A97, P98, S99, T100, L101, T102, A103, E105, R111, C124, A126, T142, S143, K147, D161, N162, E163, N164, Q165, K166, T167, V168, K169, A170, E171, S172, V173, M176, S177, F178, D179, Q180, S181, T187, D188, A192, V195, V222, N230, A232, K245, F247, D250, L251, T252, D256, A257, A258, A276, A295, A299, D300, K308, S309, A310, T311, A312, I313, F314, V316, A326, D328, K330, T331, G332, A333, E334, G335, Q336, L337, G338, M351, C357, T364, I365, V373, V375, T377 |  |  |  |  |
| NP_220232.1 | Outer membrane protein B | 10 | A61, C71, V91, V96, D101, T120, V125, R287, K297, V314 | V91, V96, D101, R287, K297, | None | C71, T120, V125, V314 | A61 |
| **GROUP C** |  |  |  |  |  |  |  |
| NP_219509.1 | Hypothetical protein CT_007 | 8 | F44, R65, P78, L109, D151, G160, E174, S274 | L109, D151, G160 | R65, F44, S274, P78 | E174 | None |
| NP_219858.1 | Hypothetical protein CT_351 | 16 | H15, S19, G40, D56, R59, G89, I93, G230, E286, V304, L373, R404, N431, N504, H511, F693 | N431, V304 | H511, G230, L373, R404, N504 | E286 | H15, S19, G40, D56, R59, G89, I93, G230, R693 |
| NP_219881.1 | Porin AaxA | 16 | D13, S38, C58, P65, A105, R159, A255, S256, H330, I331, F357, A360, V382, P388, T390, T403 | T390, P388, A105, | I331, S256, A255 | R159, F357, T403, A360, H330, V382 | C58, P65, D13, S38 |
| NP_219899.1 | Hypothetical protein CT_389 | 8 | A13, H42, I101, P166, L182, I203, V285, L311 | L182, L311 | I203 | V285 | A13, H42, I101, P166 |

**^≠^**Proteins were named as per their annotation in NCBI and UniProt databases, searched using Sequence Identifier.

ECL: Extracellular Loop; ICL: Intracellular Loop and TM: Transmembrane

^∞^Since amino acid sequence variations were present at 93 positions, therefore we have not determined whether these variations are present on the TM region or loop region of the predicted proteins.

**Table S5**: Top 20 conformational B-cell epitopes selected for the identified proteins.

| **Protein Accession No.** | **Residues** | **Number of Residues** | **Score** |
| --- | --- | --- | --- |
| NP_219746.1 | A:C26, A:S27, A:T28, A:S29, A:E30, A:G31, A:R32, A:M33, A:V34, A:V35, A:E36, A:S37, A:I38, A:T39, A:I40, A:T41, A:T42, A:Q43, A:G44, A:E45, A:N46, A:T47, A:Q48, A:N49, A:K50, A:R51, A:A52, A:I53, A:P54, A:K55, A:I56, A:K57, A:T58, A:K59, A:Q60, A:G61, A:T62, A:L63, A:F64, A:S65, A:K78, A:D79, A:F80, A:D81, A:R82, A:V83, A:E84, A:P85, A:I86, A:V87, A:E88, A:F89, A:R90, A:N91, A:G92, A:Q93, A:A94, A:V95, A:I96, A:S97, A:L98, A:I99, A:L100, A:T101, A:A102, A:K103, A:P104, A:V105, A:I106, A:R107, A:E108, A:I109, A:N110, A:I111, A:S112, A:G113, A:N114, A:E115, A:A116, A:I117, A:P118, A:T119, A:H120, A:K121, A:I122, A:L123, A:L126, A:L128, A:Y129, A:K130, A:N131, A:D132, A:L133, A:F134, A:D135, A:Y150, A:G154, A:Y155, A:Y156, A:D157, A:S158, A:Q159, A:S161, A:Y162, A:S163, A:H164, A:N165, A:H166, A:N167, A:E168, A:K169, A:E170, A:G171, A:F172, A:I173, A:D174, A:I175, A:S176, A:I177, A:E178, A:I179, A:K180, A:E181, A:G182, A:R183, A:H184, A:G185, A:R186, A:I187, A:K188, A:K189, A:L190, A:T191, A:I192, A:S193, A:G194, A:I195, A:T196, A:T198, A:E199, A:A200, A:Q211, A:Y212, A:S213, A:T214, A:T215, A:T216, A:G223, A:V224, A:Y225, A:H226, A:F241, A:Q242, A:N243, A:G245, A:Y246, A:A247, A:D248, A:A249, A:K250, A:V251, A:S252, A:K253, A:E254, A:V255, A:S256, A:T257, A:D258, A:A259, A:K260, A:G261, A:N262, A:I263, A:T264, A:L265, A:L266, A:I267, A:V268, A:V269, A:D270, A:K271, A:G272, A:P273, A:L274, A:Y275, A:T276, A:L277, A:G278, A:H279, A:V280, A:H281, A:I282, A:E283, A:G284, A:F285, A:T286, A:A287, A:L288, A:S289, A:K290, A:L292, A:L293, A:D294, A:K295, A:Q296, A:L297, A:L298, A:V299, A:G300, A:P301, A:N302, A:S303, A:L304, A:Y305, A:C306, A:Y321, A:G325, A:Y326, A:V327, A:N328, A:T329, A:N330, A:V331, A:D332, A:V333, A:S334, A:F335, A:S336, A:A337, A:H338, A:P339, A:T340, A:L341, A:P342, A:V343, A:Y344, A:D345, A:V346, A:T347, A:Y348, A:R349, A:V350, A:S351, A:E352, A:G353, A:S354, A:P355, A:Y356, A:K357, A:I358, A:G359, A:L360, A:F381, A:P382, A:G383, A:D384, A:T385, A:F386, A:R413, A:S414, A:Q415, A:L416, A:D417, A:P418, A:L419, A:D420, A:S421, A:N422, A:D423, A:L424, A:Y425, A:R426, A:D427 | 273 | 0.745 |
| NP_219746.1 | A:S23, A:S24, A:F25 | 3 | 0.742 |
| NP_220232.1 | A:M1, A:S2, A:S3, A:K4, A:L5, A:V6, A:N7, A:L9, A:R10, A:L11, A:T12, A:F13, A:L14, A:S15, A:F16, A:L17, A:G18, A:I19, A:A20, A:S21, A:T22, A:S23, A:L24 | 23 | 0.811 |
| NP_220140.1 | A:T233, A:L234, A:T235, A:A236, A:K237, A:T238, A:N239, A:D240, A:P241, A:A242, A:D243, A:A244, A:S245, A:A246, A:A247, A:Q248, A:P249, A:A250, A:K251, A:P252, A:N253, A:T254 | 22 | 0.85 |
| NP_220140.1 | A:G8, A:V10, A:G11, A:A12, A:L13, A:L14, A:L15, A:A16, A:C17, A:G18, A:S19, A:T20, A:N21, A:L22, A:A23, A:F24, A:A25, A:Q26, A:A27, A:S28, A:S29 | 21 | 0.779 |
| NP_220200.1 | A:L4, A:L5, A:K6, A:S7, A:V8, A:L9, A:V10, A:F11, A:A12, A:A13, A:L14, A:S15, A:S16, A:A17, A:S18 | 15 | 0.809 |
| NP_219509.1 | A:L2, A:K3, A:V4, A:L5, A:F6, A:H7, A:T8, A:T10, A:L11, A:F12 | 10 | 0.814 |
| NP_219509.1 | A:D30, A:R31, A:D32, A:K35, A:N36, A:P37, A:P38, A:L39 | 8 | 0.728 |
| NP_219858.1 | A:K2, A:R3, A:F4 | 3 | 0.958 |
| NP_219858.1 | A:A27, A:V28, A:Q29, A:K30, A:K31, A:I32, A:S33, A:Y34 | 8 | 0.933 |
| NP_219858.1 | A:R454, A:H455, A:K505, A:H506 | 4 | 0.885 |
| NP_219858.1 | A:L35, A:S36, A:H37, A:F38, A:K39, A:G40, A:I41, A:T42, A:G43, A:I44, A:M45, A:D46, A:V47, A:E48, A:D49, A:G50, A:V51, A:L52, A:H53, A:I54, A:H55, A:D56, A:D57, A:L58, A:R59, A:L60, A:Q61, A:A62, A:N63, A:K64, A:A65, A:Y66, A:V67, A:E68, A:N69, A:R70, A:T71, A:D72, A:C73, A:G74, A:I75, A:K76, A:I77, A:V78, A:A79, A:H80, A:G81, A:N82, A:V83, A:M84, A:V85, A:N86, A:Y87, A:R88, A:G89, A:I93, A:C94, A:D95, A:Y96, A:L97, A:E98, A:Y99, A:Y100, A:E101, A:D102, A:T103, A:D104, A:S105, A:C106, A:L107, A:L108, A:T109, A:N110, A:G122, A:S123, A:T124, A:T126, A:I127, A:S128, A:P129, A:S130, A:H135, A:K136, A:D154 | 84 | 0.812 |
| NP_219858.1 | A:F5, A:P6, A:L7, A:F8, A:I9, A:G10, A:V11, A:L12, A:L13, A:A14, A:H15, A:T16, A:L17, A:P18, A:S19, A:E20, A:G21, A:L22, A:S23, A:H24, A:Q25, A:Q26 | 22 | 0.774 |
| NP_219881.1 | A:E33, A:A34, A:C35, A:N36, A:T37, A:S38, A:S39, A:L40, A:S41, A:K42, A:E43, A:L44, A:I45, A:P46, A:L47, A:S48, A:E49, A:R51, A:G52, A:L53, A:L54, A:S55, A:P56, A:I57, A:C58, A:D59, A:F60, A:I61, A:S62, A:E63 | 30 | 0.911 |
| NP_219881.1 | A:D23, A:L24, A:I25, A:S26, A:P27, A:K28, A:P29, A:T30, A:E31 | 9 | 0.815 |
| NP_219881.1 | A:P65, A:C66, A:L67, A:H68, A:G69, A:V70, A:S71, A:V72, A:N74, A:L75, A:Q77, A:A78, A:L79, A:K80, A:N81, A:S82, A:A83, A:G84 | 18 | 0.785 |
| NP_219899.1 | A:E26, A:P27, A:N28, A:S29, A:C30, A:P31, A:D32, A:C33, A:N35, A:N36 | 10 | 0.984 |
| NP_219899.1 | A:K3, A:P4, A:L5 | 3 | 0.878 |
| NP_219899.1 | A:R185, A:F222, A:D223, A:L224, A:D225, A:H226, A:P227, A:E228, A:A229, A:C230, A:M231, A:V232, A:L262, A:G263, A:D264, A:E265, A:F266, A:I267, A:L268, A:A269, A:N270, A:Q271, A:L272, A:P273, A:P274, A:K275, A:K276, A:R277 | 28 | 0.823 |
| NP_219899.1 | A:F7, A:G8, A:Y9, A:F10, A:F11, A:C12, A:A13, A:I14, A:Y15, A:F16, A:T17, A:L18, A:L19, A:Q20, A:A21, A:A22, A:F23, A:A24, A:K25 | 19 | 0.772 |

**Supplemental Figures**

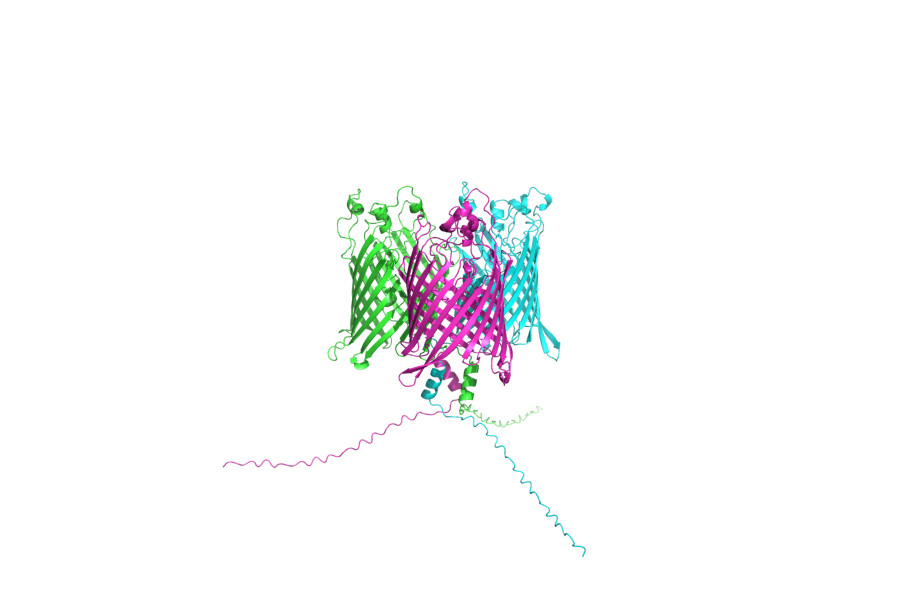
**Figure S1**: The trimeric model of NP_220140.1, generated using AlphaFold 3 exhibited strong structural confidence, with a predicted TM-score (pTM) of 0.78 and inter-chain predicted TM-score (ipTM) of 0.76, suggesting a potential homo-trimeric conformation.
